# Intrinsic and extrinsic factors regulate FtsZ function in *Caulobacter crescentus*

**DOI:** 10.1101/2023.09.08.556907

**Authors:** Jordan M Barrows, Ashley S Anderson, Barbara K Talavera-Figueroa, Erin D Goley

## Abstract

Bacterial cell division is crucial for replication and requires careful coordination via a complex set of proteins collectively known as the divisome. The tubulin-like GTPase FtsZ is the master regulator of this process and serves to recruit downstream divisome proteins and regulate their activities. Upon arrival at mid-cell, FtsZ associates with the membrane via anchoring proteins and exhibits treadmilling motion, driven by its GTP binding and hydrolysis activities. Treadmilling is proposed to play a role in Z-ring condensation, as well as in distribution and regulation of peptidoglycan (PG) cell wall remodeling enzymes. FtsZ polymer superstructure and dynamics are central to its function, yet their regulation is incompletely understood. We sought to address these gaps in knowledge by modulating intrinsic and extrinsic regulators of FtsZ and evaluating their effects *in vitro* and in cells, alone and in combination. To do this, we leveraged the cell cycle control features of *Caulobacter crescentus.* We observed that *Caulobacter* FtsZ variants that abrogate GTP hydrolysis impact FtsZ dynamics and Z-ring positioning, with little to no effect on Z-ring structure or constriction. Production of an FtsZ variant lacking its disordered C-terminal linker (ΔCTL) resulted in aberrant Z-ring dynamics and morphology, misregulated PG metabolism, and cell lysis. Combining ΔCTL and GTPase mutations was additive, suggesting they each act independently to control the Z-ring. Modulating levels of FtsA resulted in formation of multiple Z-rings that failed to constrict, suggesting roles in regulating both FtsZ superstructure and the activity of downstream divisome components. Collectively, our results indicate that GTP hydrolysis serves primarily to position the Z-ring at mid-cell, the CTL regulates both Z-ring structure and downstream signaling, and FtsA contributes to all aspects of FtsZ assembly and function. The additive effects of these elements are required to support robust and efficient cell division.

## Introduction

Cell division in bacteria is a complex process that is essential for replication. In most bacteria, cell division occurs by actively remodeling the peptidoglycan (PG) cell wall, which drives constriction of the bacterial envelope and eventually results in cell separation. In rod-shaped organisms, failure to properly regulate this process leads to filamentation and eventual lysis. Cell division is carried out by a multiprotein complex known as the “divisome,” which comprises over 20 different proteins, including PG synthases that are responsible for building the cell wall. While the identities of most players in the divisome have been known for some time, there are still fundamental questions regarding the interactions between and regulation of members of the divisome.

The tubulin homolog FtsZ is the master regulator of the divisome and is responsible for several functions prior to and throughout cell division: i) localization to mid-cell to form a “Z-ring” and mark the future site of division, ii) recruitment of downstream members of the divisome, and iii) regulation of divisome activity and distribution of PG synthases. In *Caulobacter crescentus* (hereafter *Caulobacter*), Z-ring formation is mediated by FtsZ self-assembly and its interaction with positive (ZapA/ZauP) (Woldemeskel *et al*., 2017) and negative (MipZ) (Thanbichler and Shapiro, 2006) regulators of polymerization. While ZapA/ZauP primarily affect Z-ring morphology, MipZ is responsible for Z-ring positioning, depolymerizing FtsZ at the cell poles and allowing formation of a stable Z-ring only at mid-cell.

In numerous bacteria, FtsZ has been shown to exhibit treadmilling dynamics, polymerizing and depolymerizing in a polarized manner to result in net movement of polymers along the inner circumference of the cell (Bisson-Filho *et al*., 2017; Yang *et al*., 2017). Treadmilling dynamics are linked to the rate of GTP hydrolysis, as mutants with decreased hydrolytic activity have decreased treadmilling rates (Bisson-Filho *et al*., 2017; Yang *et al*., 2017). In *Escherichia coli*, FtsZ treadmilling helps to distribute PG synthases about the division plane, and mutations that reduce treadmilling velocity also affect the movement rates of these enzymes (Yang *et al*., 2017, 2021).

FtsZ comprises three domains: i) the globular GTPase domain, ii) the disordered C-terminal linker (CTL), and the conserved C-terminal peptide (CTC). While the GTPase and CTC are well conserved across bacterial species, the CTL is poorly conserved in both length and sequence. Previous investigations from our group implicated the CTL as a key regulator of FtsZ function in *Caulobacter*, as cells expressing a mutant lacking the CTL (ΔCTL) form bulges at the incipient division site, reflecting misregulation of PG synthase activity (Sundararajan *et al*., 2015; Sundararajan and Goley, 2017; Barrows *et al*., 2020). While wild-type FtsZ forms GTP-dependent single polymers *in vitro*, ΔCTL forms bundles of polymers and exhibits decreased GTP turnover, indicating that the CTL is essential for modulating polymer structure and dynamics. The mechanisms underlying CTL-mediated effects on FtsZ assembly and the downstream function of the Z-ring are still largely unclear.

FtsA is a widely conserved actin homolog that serves as a membrane anchor for FtsZ, binding to the CTC peptide. Numerous studies have demonstrated the importance of this interaction in division, and FtsA has been implicated in the signaling pathway(s) from FtsZ to the PG synthetic machinery in *Escherichia coli* (Dai and Lutkenhaus, 1992; Pichoff and Lutkenhaus, 2005, 2007; Lutkenhaus *et al*., 2012; Szwedziak *et al*., 2012). FtsA is also able to remodel FtsZ structures *in vitro* (Szwedziak *et al*., 2012; Loose and Mitchison, 2014; Conti *et al*., 2018; Schoenemann *et al*., 2018) and *in vivo*, in some cases resulting in large-scale helices (Barrows *et al*., 2020), suggesting that it plays roles in regulating FtsZ dynamics and polymer structure as well. However, the effects of FtsA on FtsZ assembly and function and how those relate to divisome activity are not completely understood.

*Caulobacter* is a Gram-negative alphaproteobacterium that has long served as a model for bacterial morphogenesis and cell-cycle progression (Barrows and Goley, 2023). While the majority of divisome components are well-conserved from *Caulobacter* to other well-studied Gram-negative bacterial models, *Caulobacter* lacks the membrane anchor ZipA, which is specific to gammaproteobacteria (Hale and De Boer, 1997), and possesses FzlC and FzlA, FtsZ binding partners that are specific to alphaproteobacteria (Goley *et al*., 2010; Meier *et al*., 2016; Lariviere *et al*., 2018). Additionally, Z-ring localization is regulated by the MipZ system in *Caulobacter* (Thanbichler and Shapiro, 2006), which is not present in other organisms in which FtsZ dynamics have been studied (Barrows and Goley, 2021). *Caulobacter* cultures can also be synchronized at the swarmer stage of the cell cycle (Schrader and Shapiro, 2015), allowing for precise analysis of cell-cycle dependent events (e.g., Z-ring formation, constriction). This ability to synchronize *Caulobacter* makes it a desirable model system in which to evaluate FtsZ’s role in cell division. Although *Caulobacter* has been leveraged as a model for understanding cell division, a thorough study of the regulation of *Caulobacter* FtsZ structure and dynamics is lacking and is fundamental to a mechanistic understanding of cell division.

Here, we explored the roles of intrinsic and extrinsic factors on *Caulobacter* FtsZ function. By examining GTPase-deficient FtsZ and FtsZ without its CTL, we delineated the individual and combined effects of these elements. While the CTL was required for both proper Z-ring morphology and regulation of PG synthesis, GTP hydrolysis was primarily responsible for proper Z-ring placement. Finally, we found that FtsA modulates the structure and stability of FtsZ polymers in a manner that is important for downstream divisome functions. Surprisingly, the effects of these elements were largely independent of each other, resulting in additive phenotypes when combined. Overall, our results support a model wherein GTP turnover, the presence of the CTL, and FtsZ interaction with FtsA contribute to FtsZ’s function independently, forming a multifaceted control network that is required for precise regulation of PG remodeling to ensure successful cell division.

## Results

### A GTPase variant of FtsZ is deficient for GTP turnover but not polymerization

To begin assessing the role of GTPase activity in the function of *Caulobacter* FtsZ, we made two mutations based on previously studied variants: G109S and D216A (G105S and D212A in *E. coli*, respectively). These substitutions each result in a decrease in GTP turnover in *E. coli*. However, the G105S mutant is able to complement loss of wild-type *ftsZ* while the D212A mutant is not (Stricker and Erickson, 2003), suggesting that they have differing effects on FtsZ dynamics and function. We introduced these mutations into both full-length and ΔCTL FtsZ from *Caulobacter* in an effort to understand the individual and combined contributions of the CTL and GTPase activity to FtsZ function. We began our investigation by quantifying GTP turnover rates of full-length and ΔCTL *Caulobacter* FtsZ. Wild-type FtsZ and ΔCTL exhibited GTP turnover rates comparable to what we observed previously (Figure 1A and Supplemental Figure 1A) (Sundararajan and Goley, 2017). Consistent with observations in *E. coli*, G109S and D216A variants of full-length or ΔCTL FtsZ exhibited negligible GTP turnover (Figure 1A and Supplemental Figure 1A), indicating that these variants are deficient for binding or hydrolyzing GTP, or both.

**Figure 1:**
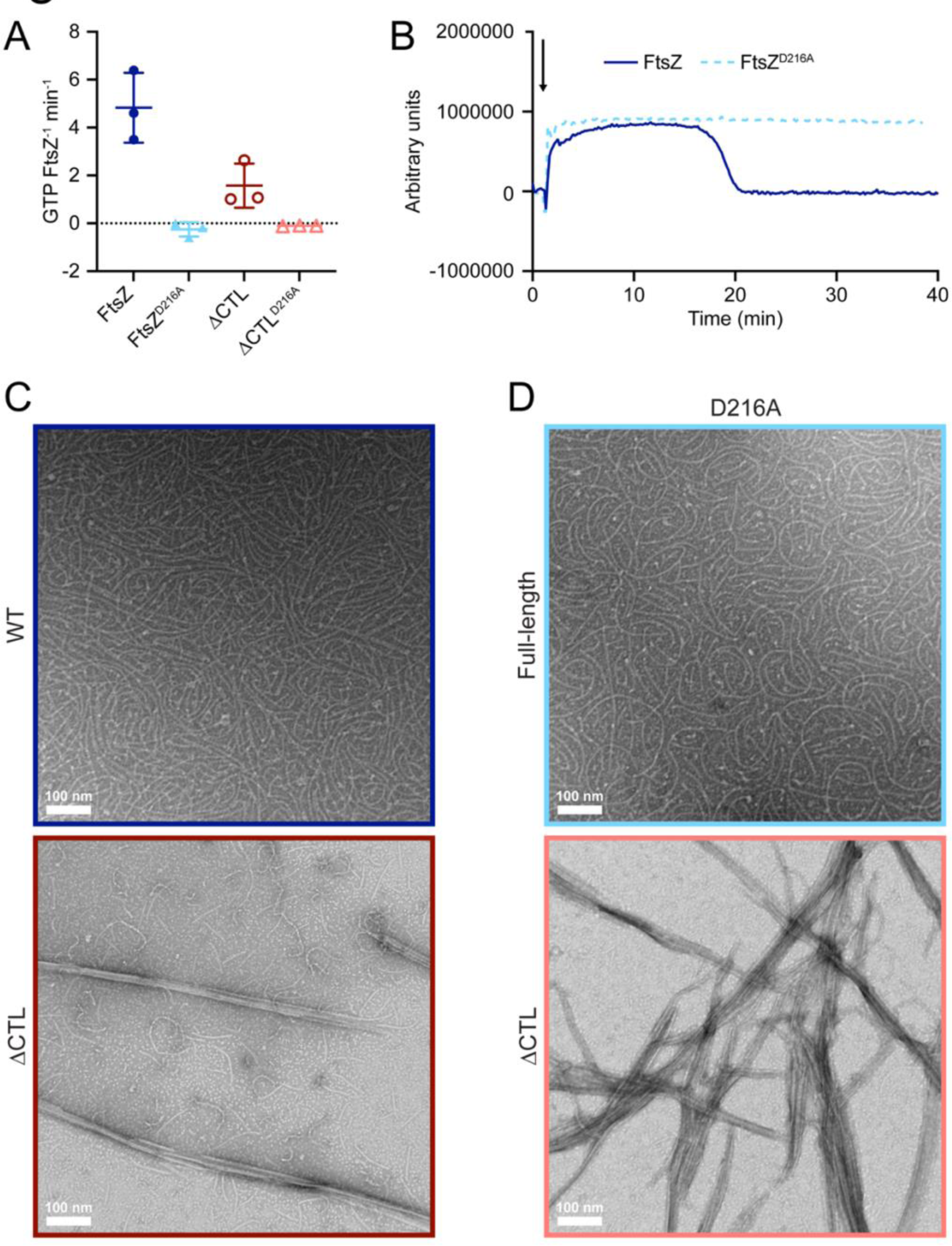
D216A variants of FtsZ exhibit decreased GTP turnover and dynamics. **A.** GTP turnover rates for indicated FtsZ variants (4 µM). **B.** Right-angle light scattering at 350 nm over time for 4 μM of each indicated protein upon addition of 0.5 mM GTP, indicated by the black arrow. Representative lines of two independent replicates are shown. **C.-D.** Representative TEM micrographs for 2 μM of each indicated protein incubated with 2 mM GTP.

We next employed right-angle light scattering and transmission electron microscopy (TEM) to explore the polymerization characteristics of these variants. Wild-type FtsZ exhibited an increase in light scattering upon addition of a limiting concentration of GTP, indicative of polymer formation, followed by a return to baseline as GTP was depleted (Figure 1B). FtsZ^D216A^ formed polymers as well, which were maintained through the duration of the experiment even after wild-type FtsZ had depolymerized (Figure 1B). Both wild-type and ΔCTL FtsZ formed polymers upon addition of GTP as observed by TEM, with ΔCTL additionally forming higher-order bundles, similar to previous observations (Figure 1C) (Sundararajan and Goley, 2017; Barrows *et al*., 2020). FtsZ^D216A^ formed polymers morphologically similar to those formed by wild-type FtsZ (Figure 1D), suggesting that its ability to bind GTP and polymerize is relatively unimpaired. Additionally, like ΔCTL, ΔCTL^D216A^ formed bundled polymers as observed by TEM (Figure 1D).

In contrast to the D216A variant, we observed no light scattering for either of the G109S variants following addition of GTP (Supplemental Figure 1B), suggesting that they fail to form polymers. This finding is supported by observations via TEM, wherein we were unable to detect polymers (Supplemental Figure 1C). This is consistent with previous reports of the corresponding variant in *E. coli* FtsZ failing to form polymers or even bind to GTP *in vitro* (RayChaudhuri and Park, 1992). Thus, we are unable to recapitulate the polymer-forming activity of the G109S variant *in vitro*, precluding our ability to draw conclusions about its effect on FtsZ function. It is unlikely that this mutation confers a loss of polymerization *in vivo*, however, as this mutant can fully complement loss of wild-type *ftsZ* in *E. coli* (Stricker and Erickson, 2003). Taken together, these data demonstrate that the D216A full-length and ΔCTL variants are both competent for polymerization (and bundling, in the case of ΔCTL) but fail to hydrolyze GTP, resulting in polymer stabilization *in vitro*.

### GTPase activity is required for growth and viability in Caulobacter

Previous work with the G109S mutant indicated that production of this species in *Caulobacter* is dominant lethal with drastic morphological defects (Wang *et al*., 2001). However, in that and related studies (Li *et al*., 2007; Goley *et al*., 2010), the G109S mutant was overexpressed from a high copy replicating plasmid. Upon expression of *ftsZ^G109S^*from an integrating plasmid, either in a merodiploid or depletion strain, cells failed to form the expected “dumbbell” morphology exhibited in the overexpression strain (Supplemental Figure 2), suggesting that the dominant lethality observed previously is dose dependent. However, we were also unable to complement loss of native *ftsZ* with the G109S mutant, suggesting that unlike in *E. coli*, the G109S mutant is lethal in the absence of wild-type FtsZ. As a result of this finding and the fact that we were unable to observe polymerization of the G109S variant *in vitro*, we elected to continue this study with only the D216A mutant.

To observe the effects of GTPase deficiency *in vivo*, we constructed strains for the simultaneous depletion of wild-type FtsZ (expression driven by a vanillate-inducible promoter) and production of a desired variant under the control of a xylose-inducible promoter (Supplemental Figure 3A). This approach allowed us to cultivate these strains in the presence of wild-type FtsZ and observe the effects of substituting it with a desired variant over time. Depletion of wild-type FtsZ in liquid culture resulted in continued increase in cell density, likely as a result of filamentation, until eventual growth arrest and lysis (Supplemental Figure 3B, left). Production of FtsZ^D216A^ resulted in a reduced growth rate and final culture density, whereas both ΔCTL variants were toxic, resulting in rapid growth arrest and lysis (Supplemental Figure 3B, right). The similar lysis kinetics of ΔCTL and ΔCTL^D216A^ suggests that the effects of GTPase deficiency is overshadowed by the toxicity of ΔCTL. Finally, none of the FtsZ variants supported growth on solid media (xylose, right), suggesting that loss of the CTL and/or GTPase deficiency is lethal (Supplemental Figure 3C).

### FtsZ GTPase activity is required for proper Z-ring placement and constriction localization

To understand the function of FtsZ GTPase activity *in vivo*, we next imaged cells depleted of FtsZ and expressing either *ftsZ* or *ftsZ^D216A^* from a xylose-inducible integrated plasmid (same strains used for results in Supplemental Figure 3). Following removal of vanillate from and addition of xylose to the growth medium, FtsZ and FtsZ^D216A^ were produced to similar levels (Supplemental Figure 4). Cells of both strains continued to divide, suggesting that the divisome is functional, but over time the D216A mutant exhibited constriction localization defects (Figure 2A, aberrant constriction localization indicated by red arrowhead). This defect resulted in an increase in the mean cell length (Figure 2B) and no discernable change in maximum width (Figure 2C) by 3 hours of expression. It is important to note that while the mean length did increase, there was also a significant population of short cells (Figure 2A, B, 3-hour time point) suggesting an overall defect in length regulation. To further characterize this defect, we plotted constriction location relative to the length of the cell (Figure 2D-E), defining the pole closest to the constriction as the right-hand pole in Figure 2D. Cells producing FtsZ (navy blue) exhibited consistent constriction localization at ∼5% away from mid-cell. However, cells producing FtsZ^D216A^ had constrictions over a much greater range of the cell length, formed multiple constrictions in some cells (Figure 2D, cells with localizations with negative x-values contain at least 2 constrictions), and had constrictions farther from mid-cell on average (Figure 2E).

**Figure 2:**
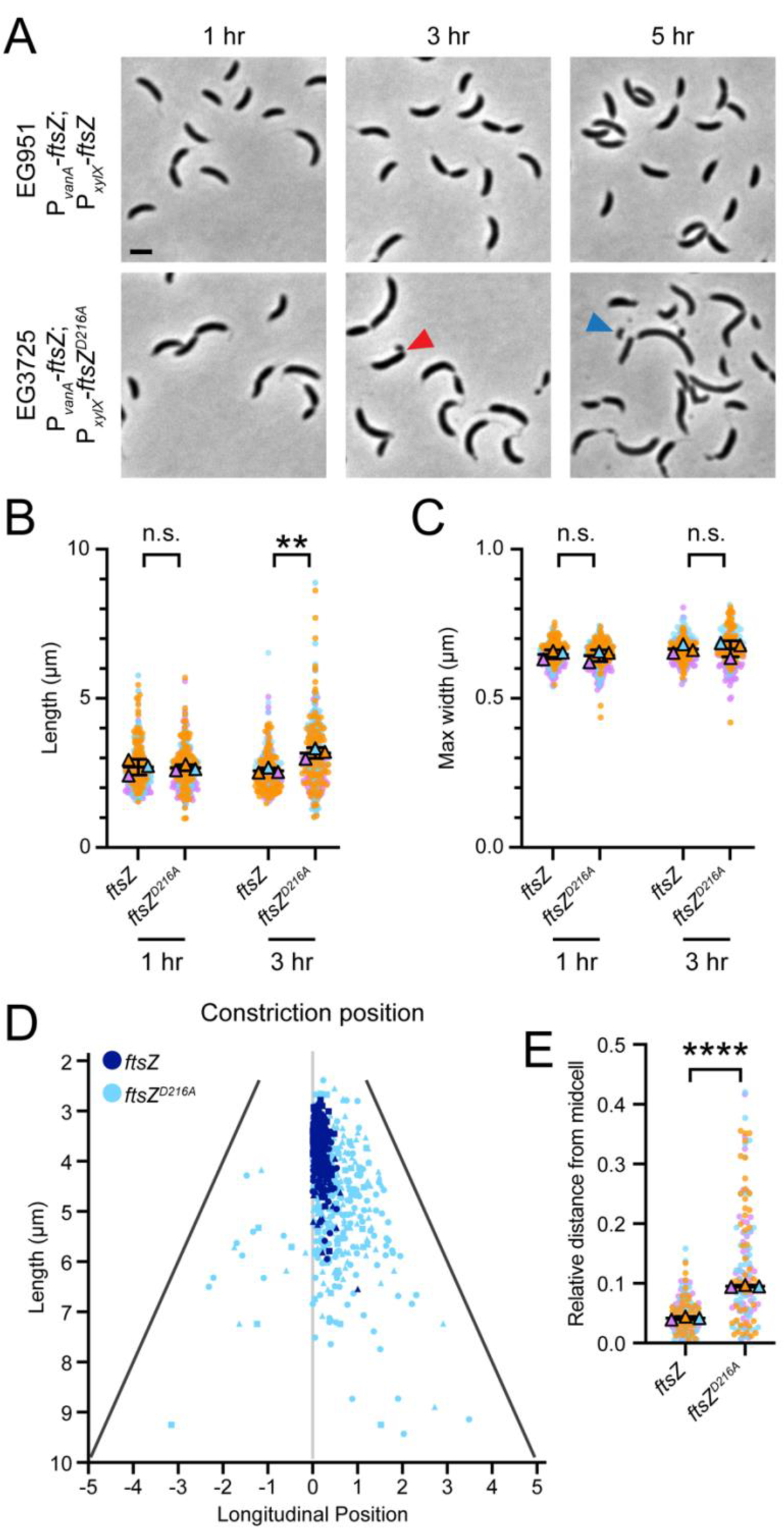
FtsZ^D216A^ production results in aberrant constriction localization. **A.** Representative phase contrast micrographs of indicated strains at given time points following simultaneous removal of vanillate and induction of xylose-driven expression of indicated *ftsZ* with 0.3% xylose. Red arrowhead indicates constriction located near the pole. Blue arrowhead indicates a minicell resulting from mislocalized constriction. Scale bar, 2 μm. B.-C. Dot plots of cell length (B) or maximum width (C) for indicated strains at 1 or 3 hrs post depletion/induction. Circles represent individual cell measurements from three independent replicates (orange, cyan, and magenta) and outlined triangles represent mean values for each replicate. Line indicates mean of replicate means and error bars are standard deviation. Unpaired t-tests were performed to determine indicated *p* values (n.s., not significant; **, *p* ≤ 0.01). D. Scatter plot of constriction position in cells from (A) at 3 hrs post depletion/induction sorted by length. The right-hand pole is defined as the pole closest to the constriction site (swarmer pole). Cells expressing *ftsZ* or *ftsZ^D216A^* are shown in dark blue and cyan, respectively. Data was acquired in three independent replicates, represented for each strain by circles, triangles, and squares. E. Dot plots of relative distance of constrictions from mid-cell in cells from (D). Circles represent individual cell measurements from three independent replicates (orange, cyan, and magenta) and outlined triangles represent median values for each replicate. Line indicates mean of replicate medians and error bars are standard deviation. An unpaired t-test was performed to determine indicated *p* value (****, *p* ≤ 0.0001).

We hypothesized that the mislocalization of constriction sites in the *ftsZ^D216A^*strain was a result of Z-ring mislocalization, which we tested by observing localization of a C-terminal fusion of mNeonGreen (mNG) to ZapA (ZapA-mNG), which binds directly to FtsZ and reports on its localization (Woldemeskel *et al*., 2017) (Figure 3). While ZapA-mNG consistently localized at mid-cell with a single focus in cells producing FtsZ, ZapA-mNG exhibited multiple bands and non-mid-cell localization in the majority of cells producing FtsZ^D216A^ (Figure 3A,B). ZapA-mNG consistently localized to rings ∼5% away from mid-cell in the wild-type strain, but ring localization was more widely distributed along the length of the cell in the D216A mutant (Figure 3B,C), reminiscent of the constriction defect we observed (Figure 2D).

**Figure 3:**
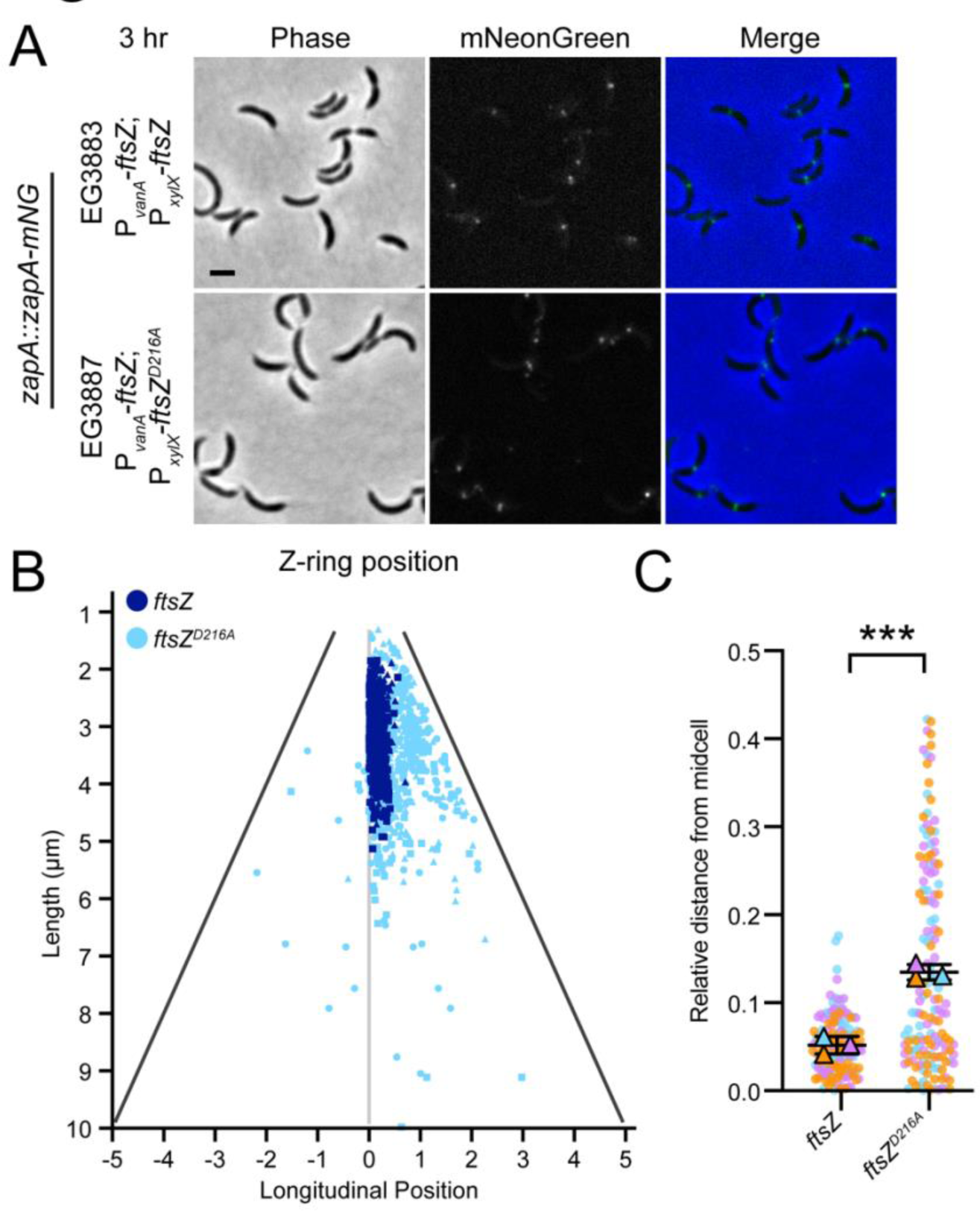
FtsZ^D216A^ production results in aberrant Z-ring localization. **A.** Representative phase contrast, epifluorescence, and merged micrographs of indicated strains with *zapA-mNeonGreen* under the *zapA* promoter at given time points following simultaneous depletion of FtsZ and induction of xylose-driven expression of indicated *ftsZ* with 0.3% xylose. Scale bar, 2 μm. **B.** Scatter plot of Z-ring position in cells from (**A**) at 3 hrs post depletion/induction sorted by length. The right-hand pole is defined as the pole closest to the Z-ring (swarmer pole). Cells expressing *ftsZ* or *ftsZ^D216A^* are shown in dark blue and cyan, respectively. Data was acquired in three independent replicates, represented for each strain by circles, triangles, and squares. **C.** Dot plots of relative distance of Z-rings from mid-cell in cells from (**B**). Circles represent individual cell measurements from three independent replicates (orange, cyan, and magenta) and outlined triangles represent median values for each replicate. Line indicates mean of replicate medians and error bars are standard deviation. An unpaired t-test was performed to determine the indicated *p* value (***, *p* ≤ 0.001).

The relationship between FtsZ dynamics and PG synthesis has been explored in *E. coli* (Yang *et al*., 2017), *Bacillus subtilis* (Bisson-Filho *et al*., 2017), *Streptococcus pneumoniae* (Perez *et al*., 2019), and *Staphylococcus aureus* (Monteiro *et al*., 2018), with differing degrees of coupling – or not - between the two depending on the species. We therefore sought to investigate whether a loss of GTPase activity in *Caulobacter* affects the kinetics of constriction, which depends on PG synthesis. Using the same strains as in Figure 2, we synchronized cultures to isolate swarmer cells, spotted them on agarose pads with growth media and vanillate, and observed their morphology over the course of a cell cycle via timelapse imaging. We used these data to measure the rates of elongation and constriction, as well as the time prior to constriction and constriction duration. As before, cells producing FtsZ^D216A^ frequently exhibited mis- localized constriction sites (Supplemental Figure 5A). However, none of the parameters we measured (elongation rate, constriction rate, elongation/constriction rate ratio, pre-constriction time, or constriction duration) were different between strains producing FtsZ or FtsZ^D216A^. We conclude that while division site placement depends on GTPase activity, constriction initiation, rate, and duration do not.

### GTPase activity is required for Z-ring positioning in the absence of the CTL

Having investigated the role of GTPase activity in full-length FtsZ, we next endeavored to examine its contribution to the phenotype resulting from production of ΔCTL to assess if GTPase activity primarily drives localization in this context as well as in the full-length version. We began by observing morphology of cells depleted of FtsZ and producing either ΔCTL or ΔCTL^D216A^ (Figure 4). As in the case of the full-length variants, both ΔCTL variants were produced in a time-dependent manner to comparable levels (Supplemental Figure 6A-B), and FtsZ was depleted to about 50% of pre-depletion levels in these strains after 1 hr (Supplemental Figure 6C). Finally, the ratio of ΔCTL to full-length FtsZ was ∼1:1 at both 3 and 5 hrs following depletion/induction (Supplemental Figure 6D). As observed previously, production of ΔCTL resulted in cell filamentation and lysis as well as bulges at the incipient division site (Figure 4A) (Sundararajan *et al*., 2015; Barrows *et al*., 2020). Production of ΔCTL^D216A^ also led to the formation of bulges, but they were smaller and more numerous, and were distributed along the length of the cell compared to larger, more centralized bulges in the *ΔCTL* strain (Figure 4A). Both strains exhibited similar increases in cell length (indicative of filamentation) and maximum width (indicative of bulging) compared to the strain expressing *ftsZ* (Figure 4B, C). To better understand the difference between these two strains, we observed the localization of the ΔCTL variants using mNG fusions (Figure 5). mNG-ΔCTL largely localized in amorphous structures at sites of bulging (Sundararajan *et al*., 2015; Barrows *et al*., 2020) (Figure 5A). mNG-ΔCTL^D216A^ adopted a more punctate localization pattern, even at early time points, with foci frequently co-localizing with bulges (Figure 5B). Thus, loss of GTPase activity hinders FtsZ’s ability to properly localize to mid-cell and distrubutes the toxic effects conferred by the loss of the CTL along the length of the cell.

**Figure 4:**
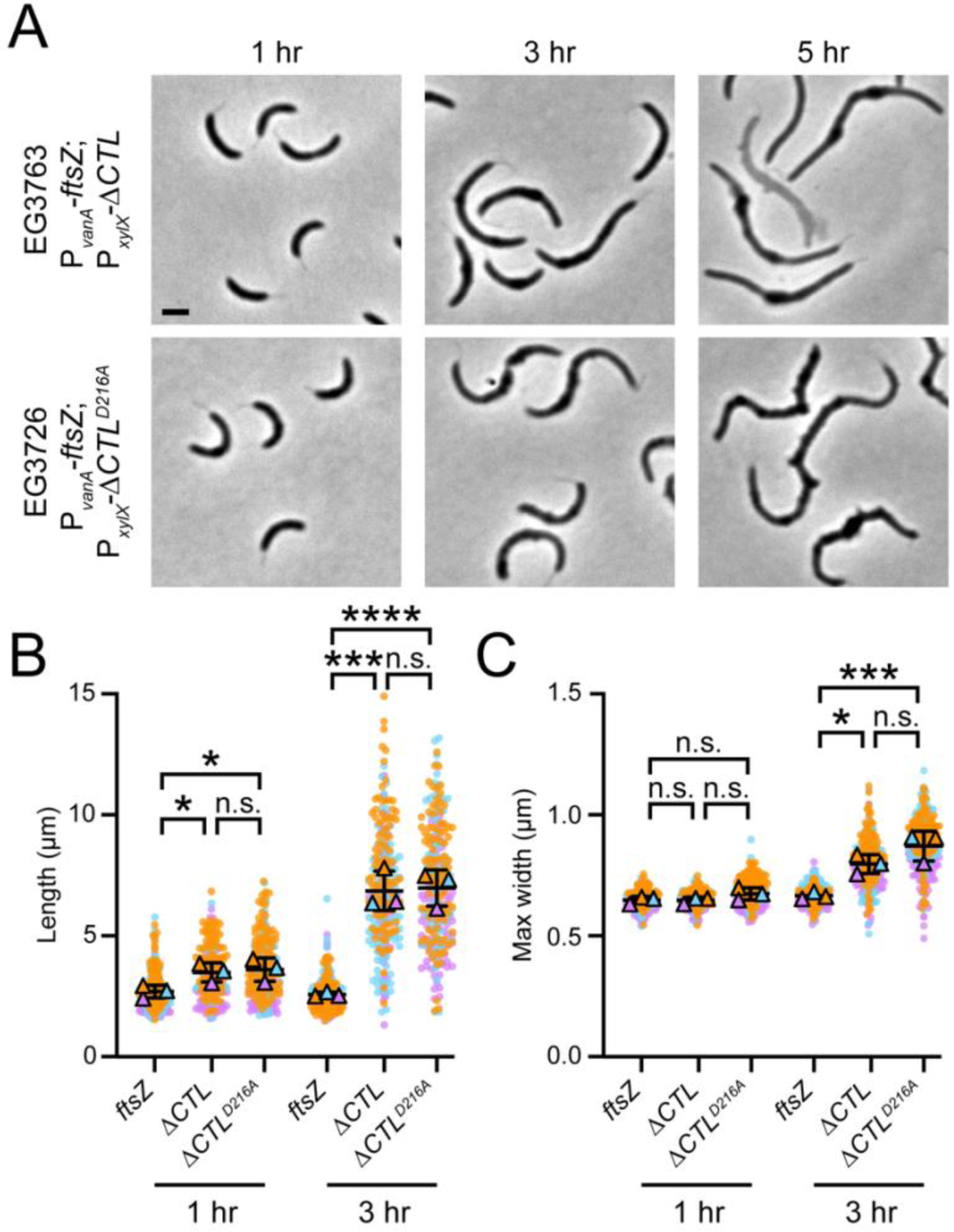
ΔCTL^D216A^ production results in bulges along the length of the cell. **A.** Representative phase contrast micrographs of indicated strains at given time points following simultaneous depletion of vanillate and induction of xylose-driven expression of indicated *ΔCTL* with 0.3% xylose. Scale bar, 2 μm. **B.-C.** Dot plots of cell length (**B**) or maximum width (**C**) for indicated strain at 1 or 3 hrs post depletion/induction. Circles represent individual cell measurements from three independent replicates (orange, cyan, and magenta) and outlined triangles represent mean values for each replicate. Line indicates mean of replicate means and error bars are standard deviation. One-way ANOVA with Šídák’s multiple comparisons test was performed to determine indicated *p* values (n.s., not significant; *, *p* ≤ 0.05; ***, *p* ≤ 0. 001; ****, *p* ≤ 0.0001).

**Figure 5:**
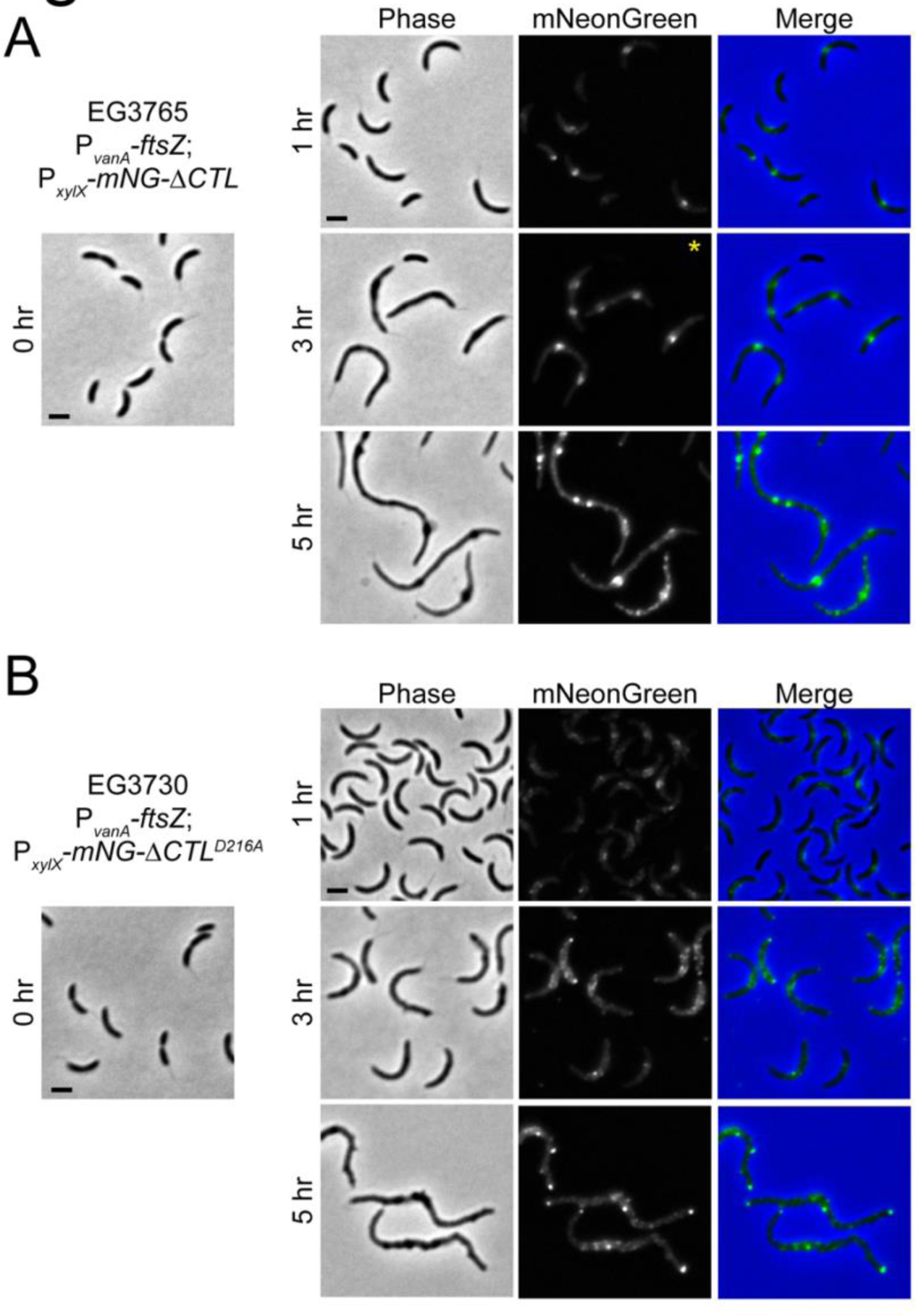
mNG-ΔCTL^D216A^ has punctate localization along the length of the cell. **A.-B.** Representative phase contrast, epifluorescence, and merged micrographs of indicated strains at given time points following simultaneous removal of vanillate and induction of xylose-driven expression of indicated *mNeonGreen-ΔCTL* fusion with 0.3% xylose. Scale bar, 2 μm. *: epifluorescence image intensity is decreased for visibility

### MipZ regulation of FtsZ depends on GTPase activity

Our data suggest that GTP hydrolysis by FtsZ is important for positioning the Z-ring at mid-cell. This finding is in line with the model proposed by (Thanbichler and Shapiro, 2006), whereby the ATPase MipZ actively depolymerizes FtsZ by stimulating its GTPase activity. As MipZ concentration is highest at the cell poles, stable polymer condensation only occurs at mid-cell where MipZ is absent. We hypothesized, therefore, that GTPase-deficient FtsZ might be insensitive to MipZ’s positioning signals. To investigate further, we expressed *mipZ* labeled with mCherry (mChy) in cells producing mNG fusions of the FtsZ variants. We observed that MipZ-mChy localized at the poles in most cells and in periodic foci corresponding to multiple origins of replication in filamentous cells regardless of the FtsZ variants present (Supplemental Figure 7), suggesting that its localization is not affected by the FtsZ variants. However, we observed multiple instances of mNG-FtsZ^D216A^, mNG-ΔCTL, and mNG-ΔCTL^D216A^ rings/foci that appeared close to MipZ foci or at the poles (see white arrowheads in 3-hr time point), suggesting that MipZ has decreased influence on the *in vivo* localization of these variants.

Next, we performed an FtsZ pelleting assay to determine whether the depolymerizing effect of MipZ *in vitro* (Thanbichler and Shapiro, 2006) would be mitigated by the loss of FtsZ GTPase activity (Supplemental Figure 8). As expected, the addition of MipZ resulted in a decrease in FtsZ in the pellet, indicative of depolymerization activity (Supplemental Figure 8A). However, there was no difference in the fraction of FtsZ^D216A^ in the pellet in presence versus absence of MipZ (Supplemental Figure 8A), while MipZ itself remained mostly soluble regardless of the species of FtsZ (Supplemental Figure 8B). We next tested whether we could see a decrease in polymers via TEM under the same conditions. While MipZ did affect FtsZ polymers in both cases, FtsZ^D216A^ polymers appear to be more robust upon addition of MipZ compared to wild-type FtsZ polymers, which were shorter and more highly curved than those of FtsZ^D216A^ with MipZ (Supplemental Figure 8C). Taken together, our results indicate that while FtsZ^D216A^ is not completely blind to regulation by MipZ, the loss of GTPase activity prevents depolymerization by MipZ enough to result in failure to localize to mid-cell, yielding the mis-localized constrictions and Z- rings observed (Figures 2 and 3).

### GTPase activity and the CTL contribute to FtsZ polymer dynamics

Previous reports in *E. coli* have demonstrated that FtsZ variants with attenuated GTPase activity exhibit decreased FtsZ polymer dynamics (Yang *et al*., 2017). Given our observations that both GTPase deficiency and loss of the CTL result in decreased polymer dynamics *in vitro* (Figure 1) and Z-rings with aberrant shape and/or localization *in vivo*, we hypothesized that polymer dynamics are likely disrupted *in vivo* as well. To evaluate polymer dynamics *in vivo*, we first attempted to measure mNG-FtsZ cluster velocity via fluorescence TIRF microscopy, as had been done previously in *E. coli* (Yang *et al*., 2017) and *B. subtilis* (Bisson-Filho *et al*., 2017). Due to the relatively small size of *Caulobacter*, however, we were unable to resolve individual clusters sufficiently to confidently report on velocity. Instead, we investigated polymer dynamics by measuring FtsZ monomer lifetime – that is, the length of time that a given FtsZ monomer is present in a polymer. A previous study in *B. subtilis* demonstrated that FtsZ monomer lifetime is affected by perturbations to FtsZ treadmilling dynamics (Squyres *et al*., 2021), so we reasoned that the variants investigated here would affect monomer lifetime as well. Indeed, imaging single molecules of FtsZ fused with HaloTag (Halo-FtsZ) (Figure 6A) and tracking the fluorescence intensity over time yielded a step-like pattern indicative of molecules that are cycling between diffusive (low intensity; monomeric) and immobile (high intensity; polymeric) states (Figure 6B), allowing us to measure the duration of each period within the polymer. Relative to wild-type FtsZ (12.2 ± 0.3 s), either the loss of the CTL (14.3 ± 0.6 s), D216A mutation (16.2 ± 0.5 s), or both (15.3 ± 0.6 s) resulted in an increase in monomer lifetime (Figure 6C), indicating either an increase in polymer stability or length, which is consistent with our prediction that these perturbations slow polymer dynamics. The differences observed in monomer lifetime were smaller than expected based on measurements in *B. subtilis* (Squyres *et al*., 2021), but this could be a result of residual wild-type FtsZ still being present at the imaging time (Figure 6D). Overall, our results are consistent with a model wherein GTPase activity and the CTL are both necessary for and modulate bulk Z-ring placement and morphology as well as individual polymer dynamics *in vivo*.

**Figure 6:**
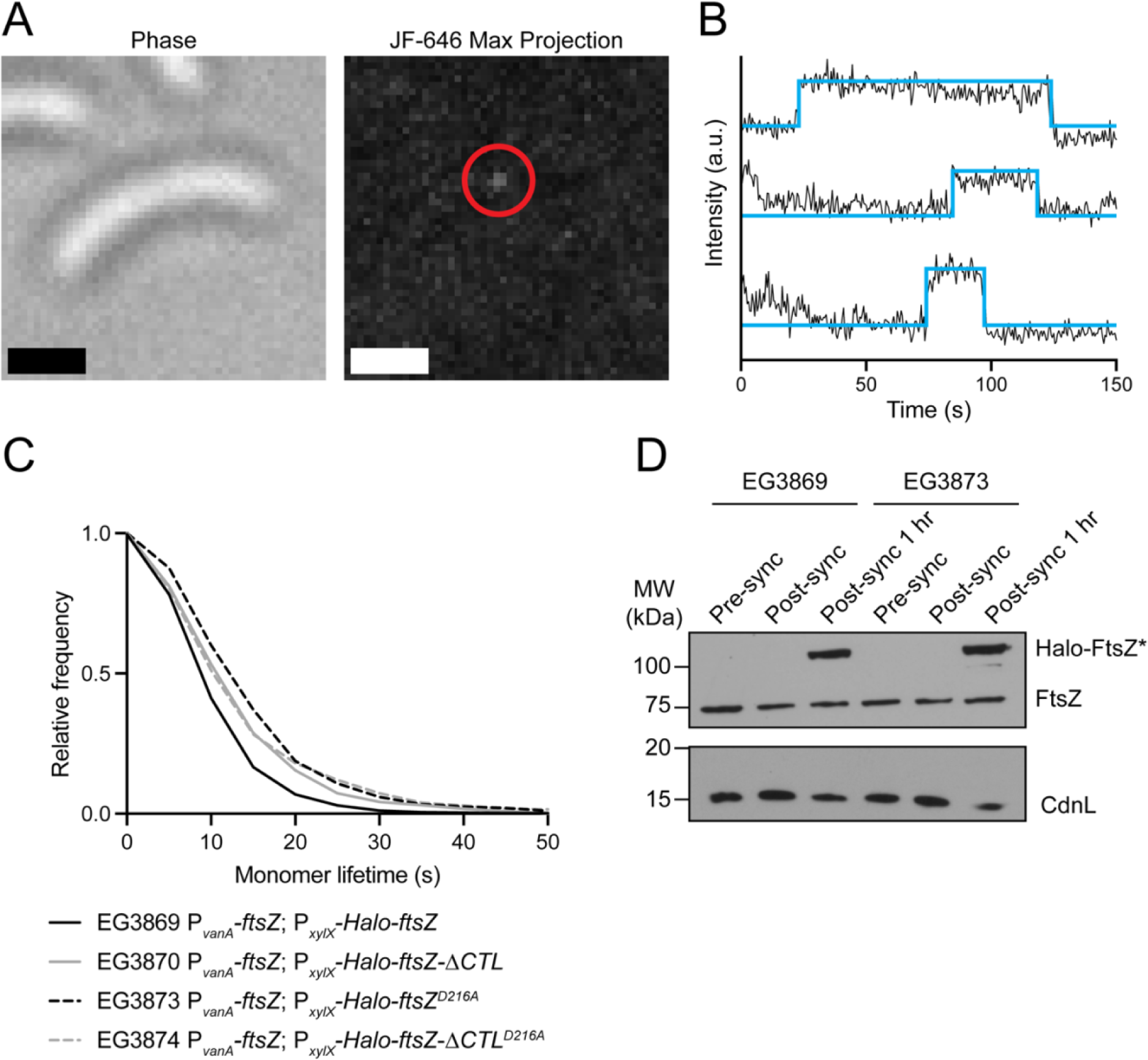
GTPase activity and the CTL modulate FtsZ polymer dynamics *in vivo*. **A.** Representative phase contrast and maximum projection of JF-646 fluorescence stack acquired for single molecule tracking of FtsZ. The molecule of interest in this stack is within the red circle. Scale bar, 1 µm. **B.** Representative intensity traces (black) and approximate fits (cyan) demonstrating step-like pattern. The length of each step was used to calculate the monomer lifetime for each trace. **C.** Frequency distribution of monomer lifetime measurements for indicated strains. **D.** Immunoblots of indicated strains bearing inducible Halo-tagged full-length FtsZ variants prior to synchrony (pre-sync), post-synchrony (post-sync), and 1 hr post-synchrony (post-sync 1 hr) to evaluate degradation of wild-type FtsZ. Blots are probed with α-FtsZ (top) and α-CdnL (bottom, loading control) primary antibodies.

### FtsA overproduction causes filamentation but only induces FtsZ helices in combination with ΔCTL

In addition to the intrinsic regulation of Z-ring positioning and function, we also wanted to examine an extrinsic factor, specifically the influence of varying levels of FtsA. We previously showed that FtsA overproduction results in filamentation and the formation of multiple Z-rings (Barrows *et al*., 2020) (Figure 7A). We next asked if these effects require GTP hydrolysis by FtsZ. The combination of mNG- FtsZ^D216A^ production and overabundant FtsA resulted in filamentous cells that form multiple Z-rings, as in the case of both the GTPase variant alone (Figure 3B) and FtsA overexpression (Figure 7A). However, unlike the case of FtsA overproduction with mNG-FtsZ, mNG-FtsZ^D216A^ was tightly clustered in only a few foci per cell when FtsA was overproduced (Figure 7B). At later time points, cells also exhibited multiple ectopic poles at sites of mNG-FtsZ^D216A^ localization.

**Figure 7:**
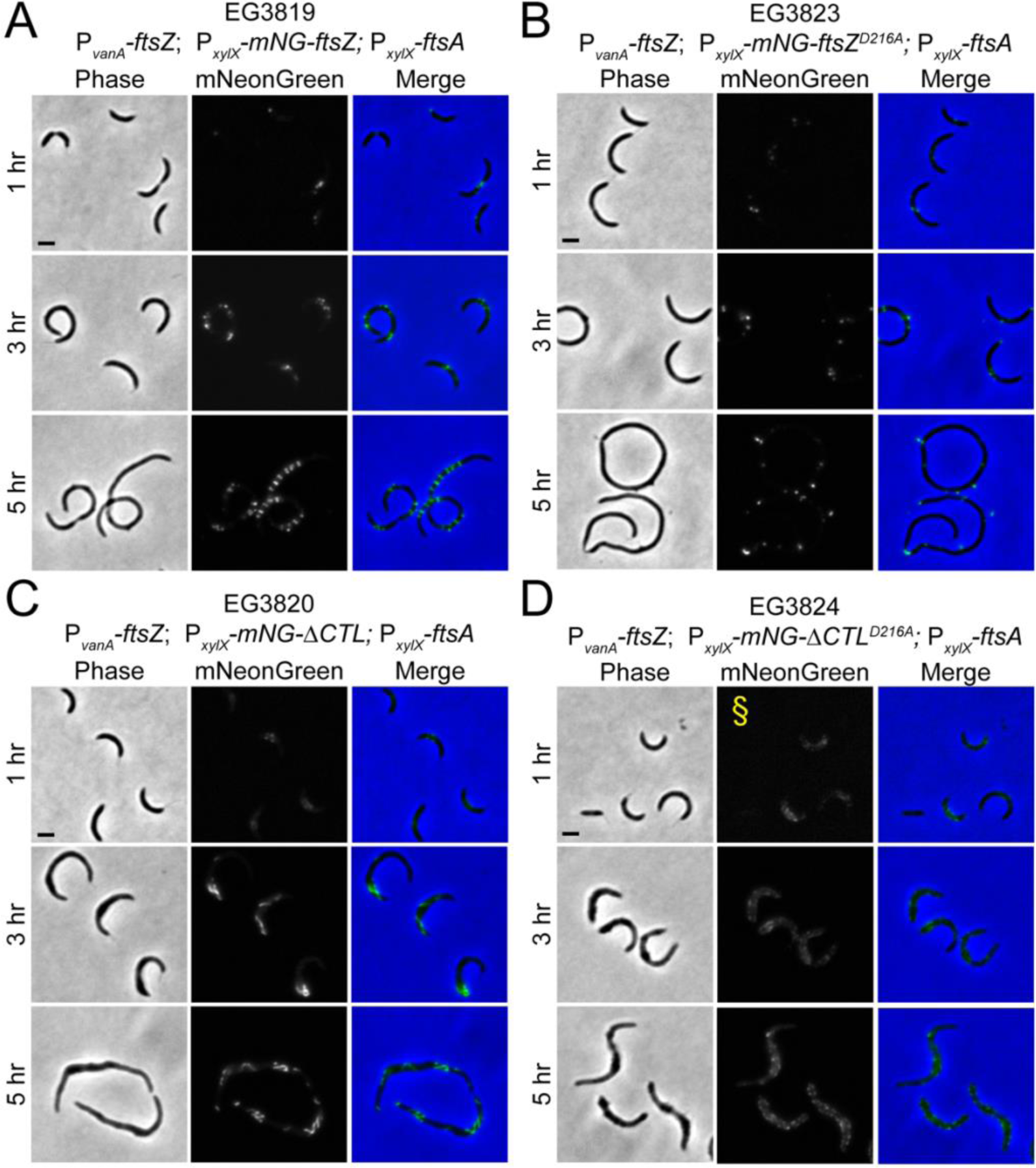
FtsA overproduction has minimal effects on FtsZ^D216A^ and ΔCTL^D216A^ phenotypes. **A.-D.** Representative phase contrast, epifluorescence, and merged micrographs of indicated strains at given time points following simultaneous depletion of vanillate and induction of xylose-driven expression of indicated *mNeonGreen-ftsZ* (**A-B**) or *mNeonGreen-ΔCTL* (**C-D**) fusion and *ftsA* with 0.3% xylose. Scale bar, 2 μm. §: epifluorescence image intensity is increased for visibility.

Overproduction of FtsA with the ΔCTL variant provided further clarity of the relationship between FtsA and GTP hydrolysis by FtsZ. While we were able to recapitulate our previous findings that mNG-ΔCTL forms large helices upon FtsA overproduction (Barrows *et al*., 2020) (Figure 7C), the phenotype with mNG-ΔCTL^D216A^ was similar to that of producing the variant with normal FtsA levels (Figure 5B) – that is, cells were filamentous with multiple bulges along their length, and exhibited numerous small FtsZ foci along the cell length (Figure 7D). Taken together, these results indicate that while FtsA overproduction can influence FtsZ localization with or without the CTL, it is unable to significantly alter the localization of GTPase-deficient FtsZ.

### FtsZ’s CTL requires both sufficient length and charge to function

Previous investigations into the role of the CTL demonstrated that FtsZ requires its CTL for function both *in vitro* and *in vivo* and that a minimum linker length of 14 amino acids is sufficient to prevent the characteristic ΔCTL bundling *in vitro* and formation of bulges in cells (Sundararajan *et al*., 2015, 2018; Sundararajan and Goley, 2017; Barrows *et al*., 2020). The *Caulobacter* FtsZ CTL contains numerous negatively charged residues (15% of total residues in CTL), implying that electrostatic repulsion plays a role in preventing stabilization via bundle formation. We sought to dissect the individual contributions of length and charge to CTL function using the aforementioned 14-amino acid CTL (L14) and a variant of L14 in which the sole glutamate residue in this shortened linker is substituted with an alanine residue (L14^E327A^). Both variants exhibited GTPase rates of ∼2 GTP FtsZ^-1^ min^-1^ (Supplemental Figure 6A), consistent with previous findings (Sundararajan and Goley, 2017). TEM of these variants following incubation with excess GTP revealed that while the E327A variant formed bundles to a degree (Supplemental Figure 6B right, white arrow heads), they were significantly less prominent compared to those formed by ΔCTL (Sundararajan *et al*., 2015) (Figure 1C), suggesting that the contribution of the single glutamate in the L14 variant CTL is only partially responsible for its lack of bundle formation (Supplemental Figure 6B, left). The morphologies of cells producing L14 or L14^E327A^ were similar, with both exhibiting filamentation, but not bulging (Supplemental Figure 6C). However, production of L14^E327A^ resulted in poorer cell growth and earlier lysis compared to L14, but less severe lysis than ΔCTL (Supplemental Figure 6D). This intermediate phenotype suggests that length and charge are necessary for complete CTL function, but that complete loss of both is required for the drastic bundling, bulging, and lysis effects caused by ΔCTL.

## Discussion

FtsZ is a crucial factor for initiating and regulating cell division in PG-bearing bacteria as it is responsible for demarcation of the future site of division, recruitment of downstream divisome factors, and modulation of their activity. Disruption of any of these functions has profound effects on division and morphogenesis, inspiring the need to understand each of them in detail. In the present study, we sought to dissect the relative and combinatorial contributions of intrinsic and extrinsic factors regulating FtsZ polymers in *Caulobacter*. Our findings support a model wherein GTPase activity is primarily responsible for Z-ring and divisome positioning but at least partly dispensible for divisome function, the CTL is responsible for regulating FtsZ polymer morphology and dynamics and affecting downstream regulation of PG metabolism, and binding to FtsA modulates both Z-ring position and downstream divisome function (Figure 8).

**Figure 8:**
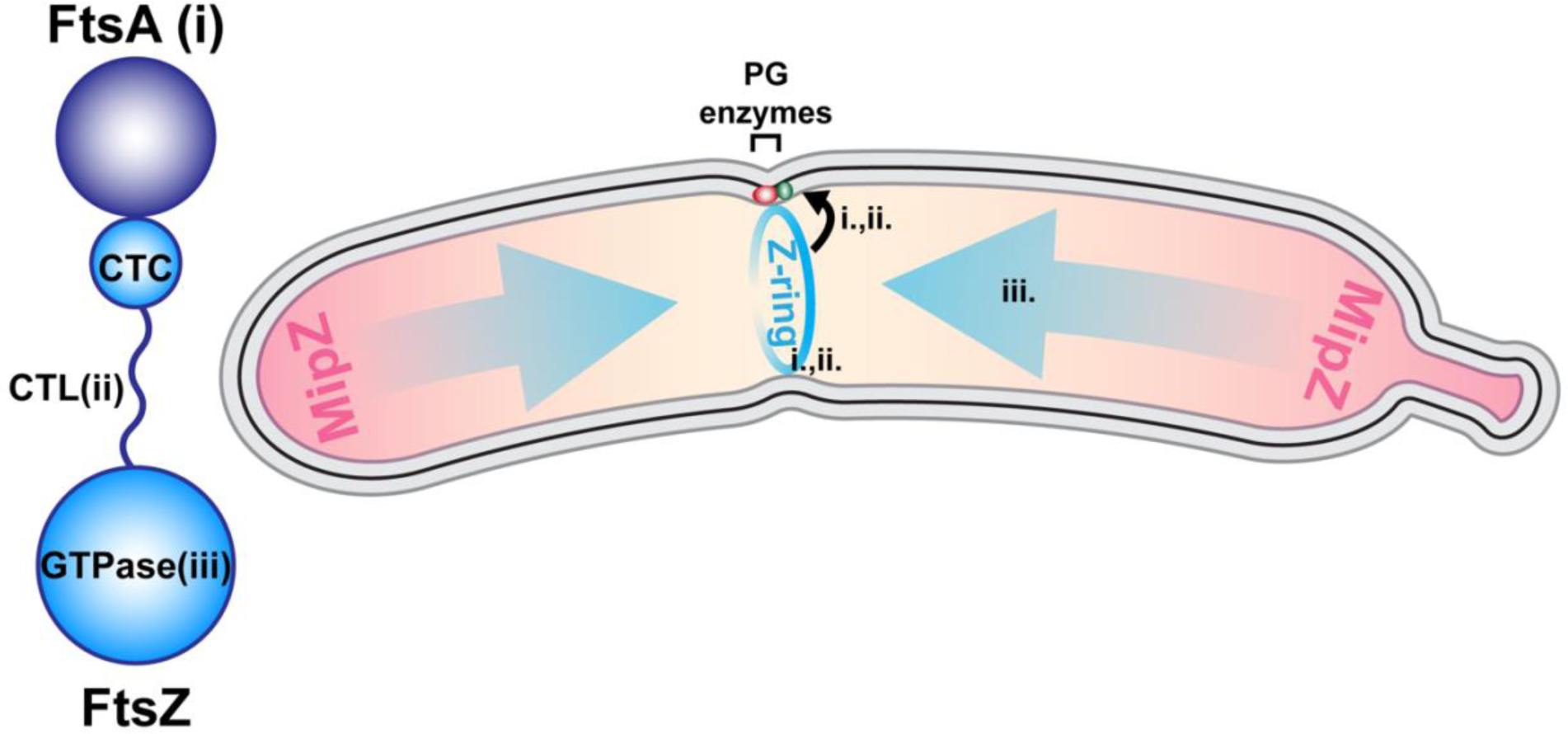
FtsA, the CTL, and GTP hydrolysis differentially affect FtsZ function. FtsZ condenses at mid-cell prior to constriction to form a Z-ring (cyan), localization of which primarily depends on regulation by MipZ (magenta) and GTP hydrolysis by FtsZ. After Z-ring formation, FtsZ recruits and regulates the activity (black curved arrow) of downstream divisome components, including PG enzymes (red and green elipses), to drive constriction of the cell envelope. FtsA levels (i.) and the CTL (ii.) both affect regulation of divisome function, with perturbation of either resulting in failure to constrict, while GTP hydrolysis (iii.) primarily regulates Z-ring localization.

Based on studies in *E. coli*, we constructed both full-length and ΔCTL variants harboring the D216A mutation, resulting in complete abrogation of GTP turnover and polymer stabilization (Figure 1).

Expression of this mutant *in vivo* causes mislocalization of Z-rings (Figure 3), resulting in a distribution of constrictions along the cell length as opposed to near mid-cell (Figure 2). Although Halo-FtsZ^D216A^ exhibits an increased dwell time in polymers *in vivo* (Figure 6), the results from our constriction rate analysis (Supplemental Figure 5) fail to indicate any defect in either constriction initation or constriction rate. Thus, GTPase activity is primarily required for Z-ring placement but is largely dispensable for Z-ring morphology and regulation of constriction. Interestingly, even though previous studies have demonstrated that PG synthase movement is correlated with FtsZ GTPase activity in *E. coli* (Yang *et al*., 2017), *Caulobacter* cells producing GTPase-deficient FtsZ are able to divide several times, although they are not viable (Supplemental Figure 3). This loss of viability suggests that regulation of Z-ring placement is crucial for propagation and that failure to properly localize the Z-ring has functional consequences, potentially resulting in aberrant segregation of cellular and genetic material due to length variation.

The mislocalization defect brought by GTPase deficiency is at least in part due to reduced responsiveness to MipZ. We observed overlapping mNG-FtsZ^D216A^ and MipZ foci in cells (Supplementary Figure 7), and MipZ had a decreased propensity to depolymerize FtsZ^D216A^ *in vitro* compared to FtsZ (Supplemental Figure 8). As MipZ has been demonstrated to enhance GTP hydrolysis by FtsZ, it makes sense that an FtsZ variant incapable of hydrolysis might be blind to regulation by MipZ, resulting in Z-ring formation even in regions containing higher MipZ levels. A more recent study suggested that MipZ depolymerizes FtsZ through multiple mechanisms (Corrales-Guerrero *et al*., 2022), but our results indicate that enhancement of GTPase activity is likely a critical form of FtsZ regulation by MipZ in cells.

GTPase activity also affects the morphology of cells producing FtsZ lacking its CTL. While the growth and viability defects of the ΔCTL^D216A^ variant mimics that of cells producing ΔCTL (Supplemental Figure 3), the former results in bulges that are smaller and more numerous, and that do not span the entire width of the cell, unlike those induced by ΔCTL (Figure 4). Additionally, mNG-ΔCTL^D216A^ localizes into several clusters, rather than a few large, amorphous Z-rings (Figure 5). This phenotype is intriguing because it suggests that the effects of ΔCTL and GTPase deficiency are modular and additive, i.e., ΔCTL results in aberrant regulation of PG metabolism, leading to bulging, while the lack of GTP hydrolysis leads to a loss of localization, resulting in more distributed foci of FtsZ along the length of the cell.

The effect of overabundant FtsA on GTPase-deficient FtsZs and the resulting phenotypes also appear to be modular. In the case of the full-length variant, FtsA overproduction results in filamentation, and cells exhibit few scattered Z-rings along their length (Figure 7), essentially a combination of the two independent phenotypes. FtsA overproduction appears to have little effect on the ΔCTL^D216A^ phenotype, as cells are morphologically similar and exhibit numerous mNG-ΔCTL^D216A^ foci. Given the fact that FtsA overproduction can drastically remodel Z-rings (Barrows *et al*., 2020) (Figure 7), this finding suggests that GTPase activity is required for modulation and/or stabilization of FtsZ polymers by FtsA. The reason for this is not clear and will be difficult to discern without accompanying biochemical data with *Caulobacter* FtsA, which we were unable to purify. Additionally, while we previously suggested that the large-scale helices formed by mNG-ΔCTL in the presence of overabundant FtsA were a result of hyperstabilization of ΔCTL by FtsA (Barrows *et al*., 2020), this hypothesis is not supported by our observation that stabilizing either FtsZ or ΔCTL by eliminating GTPase activity is not sufficient to form helices, with or without FtsA overproduction. Rather, helix formation appears to be specific to the FtsA/ΔCTL interaction.

Finally, we have expanded upon our previous investigations into the roles of the CTL in FtsZ function to discover that it requires both sufficient length and negative charge function properly (Supplemental Figure 9). This finding solidifies our hypothesis that the CTL acts as an electrostatic repulsive brush that limits lateral interactions between filaments and promotes subunit turnover (Sundararajan and Goley, 2017). It also indicates that even in the absence of negative charge, a minimal linker is not only required but is sufficient to prevent bundling and the formation of bulges. These observations support a model wherein the CTL functions to repel other FtsZ filaments, potentially serves as a buffer to prevent deleterious interactions between the GTPase domain and the CTC/FtsA/the membrane, and provides flexibility for polymers that are attached to the membrane.

Overall, our findings suggest that each of the factors investigated affects FtsZ function independently and additively: i) GTPase activity is primarily responsible for Z-ring positioning and is dispensable for constriction, ii) FtsA levels contribute to Z-ring positioning and morphology as well as downstream regulation, and iii) the FtsZ CTL is dispensable for Z-ring positioning but is crucial for proper Z-ring morphology and downstream regulation (Figure 8). Further investigation is required to understand the specific mechanism(s) by which FtsZ performs its functions and how they are influenced by both GTP hydrolytic activity and interactions with membrane anchors, such as FtsA.

## Materials and Methods

### Caulobacter crescentus *and* Escherichia coli *growth media and conditions*

*Caulobacter* NA1000 cells were grown at 30°C in peptone-yeast extract (PYE) medium. Antibiotic concentrations used in liquid (solid) media for *Caulobacter* were as follows: gentamicin, 1 (5) µg mL^-1^; kanamycin, 5 (25) µg mL^-1^; spectinomycin, 25 (100) µg mL^-1^; streptomycin, (5) µg mL^-1^. For experiments involving inducible gene expression, inducer concentrations were as follows: glucose, 0.2% (wt/vol); xylose, 0.3% (wt/vol); vanillate, 0.5 mM. *E. coli* Rosetta(DE3)/pLysS cells were grown at 37°C in Luria-Bertani (LB) medium. Antibiotic concentrations used in liquid (solid) media for *E. coli* were as follows: ampicillin, 50 (100) µg mL^-1^. Strains used in this study are listed in Supplemental Table 1. Plasmids used in this study are listed in Supplemental Table 2.

### FtsZ protein purification

Full-length and ΔCTL FtsZ, as well as GTPase variants, from *Caulobacter* were purified using the protocol described for *Cc*FtsZ in Sundararajan and Goley, 2017 (Sundararajan and Goley, 2017). *ftsZ* (or variant) expression was induced in *E. coli* Rosetta(DE3)/pLysS cells bearing *ftsZ* (or variant) on a pET21a vector. The culture was grown at 37°C to an OD_600_ of 1.0, at which point protein expression was induced using 0.5 mM isopropyl-β-D-thiogalactopyranoside (IPTG) at 37°C for 3 h. Cells were pelleted and resuspended in lysis buffer (50 mM Tris-HCl [pH 8.0], 50 mM KCl, 1 mM EDTA, 10% glycerol, DNase I, 1 mM β-mercaptoethanol, 2 mM phenylmethylsulfonyl fluoride [PMSF], 1 cOmplete mini, EDTA-free protease inhibitor tablet [Roche]) and incubated with 1 mg mL^-1^ lysozyme for 1 h at 25°C for lysis, followed by sonication. Protein was purified using an anion exchange chromatography column (HiTrap Q HP, 5 mL; GE Life Sciences) followed by ammonium sulfate precipitation (20 to 30% ammonium sulfate saturation, depending on the variant). The precipitated pellet was resuspended in FtsZ storage buffer (50 mM HEPES-KOH [pH 7.2], 50 mM KCl, 0.1 mM EDTA, 1 mM β-mercaptoethanol, 10% glycerol) and was further purified by size exclusion chromatography (Superdex 200 10/300 GL column; GE Life Sciences/Cytiva). Peak fractions were pooled, aliquoted, and snap frozen in liquid nitrogen for long-term storage at -80°C in FtsZ storage buffer.

### MipZ protein purification

His_6_-MipZ was purified as previously described (Sundararajan et al 2015). Rosetta(DE3) pLysS *E. coli* cells carrying pMT183 were grown at 37℃ to OD_600_ of 0.6 then induced with 0.5 mM IPTG for 3 h. Cells were harvested by centrifugation at 6,000 x g for 10 min at 4℃ then washed in PBS and frozen in liquid nitrogen for storage at -80℃ until purification. The cell pellet was resuspended in Ni lysis buffer (50 mM Tris-HCl [pH 8.0], 300 mM NaCl, 10 mM imidazole, 1 mM EDTA, 10% glycerol) with protease inhibitors (1 mini complete protease inhibitor table per 30 mL (Roche) and 2 mM PMSF) and 2 U/mL DNAse I. Cells were lysed by passage through a French Press at 15,000 psi. Lysates were clarified by centrifugation at 15,000 x g for 30 min at 4℃. His_6_-MipZ was purified by binding to Ni-NTA agarose (Qiagen), washed in Ni wash buffer (Ni lysis buffer with 20 mM imidazole) and eluted in Ni elution buffer (Ni lysis buffer with 300 mM imidazole). Protein was applied to a Superdex 200 10/300 GL column (Cytiva) equilibrated in 50 mM HEPES-NaOH [pH 7.2], 50 mM NaCl, 0.1 mM EDTA, 1 mM β- mercaptoethanol, and 10% glycerol. Fractions containing His_6_-MipZ were pooled, concentrated, frozen in liquid nitrogen and stored at -80℃.

### Phosphate release assay for FtsZ GTPase activity

Phosphate release by GTP hydrolysis was observed using an assay similar to that described in Sundararajan and Goley, 2017 (Sundararajan and Goley, 2017). Thawed protein was diluted to 4 µM in polymerization buffer (50 mM [300 mM for ΔCTL/L14 variants] HEPES-KOH [pH 7.2], 50 mM KCl, 0.1 mM EDTA) with 2.5 mM (10 mM for ΔCTL/L14 variants) MgCl_2_. 2 mM GTP was added and reactions were run for 0 to 30 min in 5 min intervals, stopping each reaction by adding to quench buffer (50/300 mM HEPES-KOH [pH 7.2], 50 mM KCl, 21.3 mM EDTA). Malachite green reagent (SensoLyte MG Phosphate Assay Kit [AnaSpec]) was added to each reaction and incubated for 30 min before measuring absorbance at 660 nm. Values were compared to a standard curve and plotted to determine GTPase rate. The rate for each protein was measured in triplicate.

### FtsZ polymerization kinetic assay

FtsZ polymerization kinetics were observed using an assay similar to that described in Sundararajan and Goley, 2017 (Sundararajan and Goley, 2017). Thawed protein was diluted to 4 µM in polymerization buffer (50 mM [300 mM for ΔCTL variants] HEPES-KOH [pH 7.2], 50 mM KCl, 0.1 mM EDTA) with 2.5 mM (10 mM for ΔCTL variants) MgCl_2_ in cuvettes. Following addition of a limiting concentration of GTP (0.5 mM), polymerization was measured using a Fluoromax-3 spectrofluorometer (Jobin Yvon, Inc.) to measure right-angle light scattering (excitation and emission at 350 nm, 2-nm slits) every 10 s for 40 min.

### Transmission electron microscopy (TEM)

TEM to visualize FtsZ polymers was performed as described in Sundararajan et al., 2015 (Sundararajan *et al*., 2015). Thawed protein was diluted to 2 µM in polymerization buffer (50 mM [300 mM for ΔCTL variants] HEPES-KOH [pH 7.2], 50 mM KCl, 0.1 mM EDTA) with 2.5 mM (10 mM for ΔCTL variants; 5 mM for reaction with MipZ) MgCl_2_. Polymerization was induced with addition of 2 mM GTP and reactions were incubated for 15 min prior to spotting on glow-discharged carbon-coated copper grids (Electron Microscopy Sciences, Hatfield, PA). Grids were blotted and stained twice with 0.75% uranyl formate for 2 min. Grids were then dried and imaged at 100,000 x magnification using a Hitachi 7600 TEM (operated at 80 kV) with an AMT XR80 8 megapixel CCD camera (AMT Imaging).

### High-speed pelleting assay to measure FtsZ polymerization

The fraction of steady-state polymerized FtsZ was measured similar to as described in (Goley *et al*., 2010; Sundararajan and Goley, 2017; Sundararajan *et al*., 2018). Frozen aliquots of FtsZ or FtsZ^D216A^ in storage buffer were thawed and spun at 250,000 x g for 15 min at 25°C to remove non-specific aggregates.

Clarified FtsZ was diluted to 2 µM in polymerization buffer (50 mM HEPES-KOH [pH 7.2], 50 mM KCl, 0.1 mM EDTA) with 5 mM MgCl_2_ and 2 mM ATP with or without 6 µM MipZ. Polymerization was induced with addition of 2 mM GTP, and reactions were incubated for 15 min prior to pelleting by ultracentrifugation at 250,000 x g for 15 min at 25°C. Pellet and supernatant fractions were visualized via SDS-PAGE and Coommassie staining followed by analysis using ImageLab (Bio-Rad).

### Phase-contrast and epifluorescence microscopy and image analysis

Log-phase cultures of cells were spotted onto 1% agarose pads and were imaged using a Nikon Eclipse Ti inverted microscope through a Nikon Plan Fluor 100x (numeric aperture, 1.30) oil Ph3 objective with a Photometrics CoolSNAP HQ^2^ cooled CCD camera. Background intensity was subtracted from all raw fluorescence images prior to analysis/figure preparation using the background subtraction function in FIJI (Schindelin *et al*., 2012) (rolling ball radius = 25 pixels). Images were prepared for presentation in Photoshop (Adobe) by adjusting fluorescence signal to the same levels (unless otherwise indicated) across samples in each experiment without oversaturating pixels. Length, maximum width, and constriction/Z-ring location analyses were performed using the MicrobeJ plugin for FIJI (Ducret *et al*., 2016). For the position measurements, cells were oriented using the Feature function, defining pole 1 as the pole closest to the constriction/Z-ring. Relative distance from mid-cell values were calculated by subtracting 0.5 from the arbitrary relative value generated by MicrobeJ and taking the absolute value. Each analysis was performed in biological triplicate for each strain, and comparisons were evaluated using either a student’s t-test or one-way ANOVA with Šídák’s multiple comparisons test as indicated in corresponding figure legends.

### Time-lapse imaging and constriction rate analysis

Imaging and analysis of constriction rate were performed similar to that described previously (Lariviere *et al*., 2018). Log-phase cultures of cells grown with vanillate for FtsZ production were washed with plain PYE and incubated for 30 min to begin FtsZ depletion. Cells were then synchronized (Schrader and Shapiro, 2015) and spotted onto 1% agarose pads containing PYE and xylose to induce production of the indicated FtsZ variant. Cells were then imaged by phase contrast microscopy as described above every 5 minutes until cells had completed division. Constriction/elongation rate analysis was undertaken using MicrobeJ, tracking each cell throughout the timelapse and recording the presence/absence of a constriction as well as the length and width for each frame. These values were then used to calculate the following parameters: pre-constriction/constriction time (number of frames prior to/during which a constriction is present multipled by 5 min/frame), constriction rate (cell width at constriction initiation divided by the constriction time), and elongation rate (difference of initial and final cell lengths divided by the sum of pre-constriction and constriction time).

### Cell preparation for FtsZ single molecule tracking

Log-phase cultures containing vanillate were synchronized (Schrader and Shapiro, 2015), induced with 0.3% xylose for expression of the indicated Halo-tagged mutant, and incubated at 30°C for 30 minutes. Cells were then incubated with 0.7 nM Janelia fluor 646 dye conjugated to Halo-tag ligand (JF646; Promega) in the dark at 30°C for 15 min and washed with plain PYE to remove excess dye. Cells were then mounted on a 1% agarose pad made with plain PYE and dried prior to imaging.

### Microscope and imaging for FtsZ single molecule tracking

Labeled strains were imaged on an Olympus IX-71 microscope with a 100x/1.30 NA Oil Ph3 objective and an Andor iXon 897 Ultra EM-CCD camera with EM-gain set to 300 using Metamorph software. A region of 250 x 250 pixels (160 nm/pixel) was selected, and a 300-frame image stack was acquired (0.5 s/frame) with constant illumination with a 647 nm laser set to 5 W/cm^2^. Phase contrast images were acquired before and after acquisition of fluorescence images to detect cell drift.

### Analyzing single molecule trajectories and extracting monomer lifetime values

Image stacks were analyzed in a manner similar to that described in (McCausland *et al*., 2021). Briefly, fluorescence image stacks were processed using the ThunderSTORM plugin (Ovesný *et al*., 2014) for FIJI to detect and localize particle spots. Processed data was then analyzed using custom scripts (McCausland *et al*., 2021) in Matlab R2022b to link particles to trajectories, manually unwrap cells, segment trajectories and classify trajectories, and output monomer lifetime values. Final data are the results of 3 biological replicates pooled into a single population and then presented as frequency distribution curves.

### Growth rate measurement

Log phase cultures of cells grown without inducer were diluted to an OD_600_ of 0.05 and grown in 96-well plates for 24 h at 30°C with constant shaking on a BioTek Cytation 1 plate reader (Agilent), measuring the absorbance at 600 nm every 30 min for three technical replicates for each strain. Shaded region represents standard deviation for each strain.

### Spot dilution assay

Log phase cultures of cells grown without inducer were diluted to an OD_600_ of 0.05 and serially diluted up to 10^-5^ before spotting onto solid media with indicated inducer concentrations. Plates were incubated for 48 h at 30°C and imaged with a GE Healthcare Amersham Imager 600.

### Immunoblots for measuring FtsZ levels

Indicated strains were grown to log phase in the presence of 0.5 mM vanillate, depleted of native FtsZ by washing cells in PYE, and incubated with either 0.2% glucose (control) or 0.3% xylose to induce expression of the indicated *ftsZ* mutant. Samples were collected at the indicated time points after induction and resuspended in 1x SDS loading buffer. Samples were run on 12% SDS-PAGE gels and transferred to nitrocellulose membranes, at which point they were blocked overnight at 4°C with 5% milk in TBST. Blocked blots were probed with 1° antibodies (α-FtsZ: rabbit, 1:20,000 (Sundararajan *et al*., 2015); α-CdnL: rabbit, 1:5000 (Woldemeskel *et al*., 2020)) for 1 hr at room temperature, washed 3 times for 5 minutes with TBST, probed with 2°C α-rabbit-HRP antibody (1:10,000) for 1 hr at room temperature, washed 3 more times for 5 minutes with TBST, and developed with Clarity Western ECL Substrate (Bio Rad) for subsequent imaging on a GE Healthcare Amersham 600 imager. Quantification of bands was performed in Image Lab, version 6.0 (Bio Rad), via manual band detection and automatic integration of volume. FtsZ values were adjusted with CdnL loading controls and normalized to the 0 hr time point in each replicate before being plotted using Prism (GraphPad).

## Acknowledgments

We would like to thank members of the Goley lab for helpful discussions throughout this work and feedback on the manuscript. We would also like to thank members of the Xiao lab, particularly Martin Yepes and Dr. Josh McCausland for their guidance with planning, carrying out, and performing analysis for FtsZ monomer lifetime measurements.

Work in the Goley laboratory on *Caulobacter* was supported by the National Institutes of Health (grant numbers R35GM136221 [to E.D.G.] and T32GM007445 [training grant support of and J.M.B. and B.T.F.]).

## Author contributions

J.M.B. and E.D.G. designed and performed experiments. A.S.A. performed phase and epifluorescence images of *ftsA* overexpression strains. B.K.T.F. performed phase contrast imaging for constriction rate analysis. J.M.B. carried out quantitative and statistical analysis. J.M.B. and E.D.G. conceptualized and contributed to writing and editing the manuscript.

## Abbreviations

PG: peptidoglycan
CTL: C-terminal linker of FtsZ
CTC: C-terminal conserved peptide of FtsZ
ΔCTL: FtsZ lacking C-terminal linker mChy: mCherry
mNG: mNeonGreen

## References

1. Barrows, JM, and Goley, ED (2021). FtsZ dynamics in bacterial division: What, how, and why? Curr Opin Cell Biol 68, 163–172.

2. Barrows, JM, and Goley, ED (2023). Synchronized Swarmers and Sticky Stalks: *Caulobacter crescentus* as a Model for Bacterial Cell Biology. J Bacteriol.

3. Barrows, JM, Sundararajan, K, Bhargava, A, and Goley, ED (2020). FtsA Regulates Z-Ring Morphology and Cell Wall Metabolism in an FtsZ C-Terminal Linker-Dependent Manner in *Caulobacter crescentus*. J Bacteriol 202, 1–20.

4. Bisson-Filho, AW et al. (2017). Treadmilling by FtsZ filaments drives peptidoglycan synthesis and bacterial cell division. Science (1979) 355, 739–743.

5. Conti, J, Viola, MG, and Camberg, JL (2018). FtsA reshapes membrane architecture and remodels the Z-ring in *Escherichia coli*. Mol Microbiol 107, 558–576.

6. Corrales-Guerrero, L, Steinchen, W, Ramm, B, Mücksch, J, Rosum, J, Refes, Y, Heimerl, T, Bange, G, Schwille, P, and Thanbichler, M (2022). MipZ caps the plus-end of FtsZ polymers to promote their rapid disassembly. Proceedings of the National Academy of Sciences 119.

7. Dai, K, and Lutkenhaus, J (1992). The proper ratio of FtsZ to FtsA is required for cell division to occur in *Escherichia coli*. J Bacteriol 174, 6145–6151.

8. Ducret, A, Quardokus, EM, and Brun, Y V. (2016). MicrobeJ, a tool for high throughput bacterial cell detection and quantitative analysis. Nat Microbiol 1, 16077.

9. Goley, ED, Dye, NA, Werner, JN, Gitai, Z, and Shapiro, L (2010). Imaging-Based Identification of a Critical Regulator of FtsZ Protofilament Curvature in *Caulobacter*. Mol Cell 39, 975–987.

10. Hale, CA, and De Boer, PAJ (1997). Direct binding of FtsZ to ZipA, an essential component of the septal ring structure that mediates cell division in E. coli. Cell 88, 175–185.

11. Lariviere, PJ, Szwedziak, P, Mahone, CR, Löwe, J, and Goley, ED (2018). FzlA, an essential regulator of FtsZ filament curvature, controls constriction rate during *Caulobacter* division. Mol Microbiol 107, 180– 197.

12. Li, Z, Trimble, MJ, Brun, Y V, and Jensen, GJ (2007). The structure of FtsZ filaments in vivo suggests a force-generating role in cell division. EMBO J 26, 4694–4708.

13. Loose, M, and Mitchison, TJ (2014). The bacterial cell division proteins FtsA and FtsZ self-organize into dynamic cytoskeletal patterns. Nat Cell Biol 16, 38–46.

14. Lutkenhaus, J, Pichoff, S, and Du, S (2012). Bacterial cytokinesis: From Z ring to divisome. Cytoskeleton 69, 778–790.

15. McCausland, JW et al. (2021). Treadmilling FtsZ polymers drive the directional movement of sPG-synthesis enzymes via a Brownian ratchet mechanism. Nat Commun 12, 609.

16. Meier, EL, Razavi, S, Inoue, T, and Goley, ED (2016). A novel membrane anchor for FtsZ is linked to cell wall hydrolysis in *Caulobacter crescentus*. Mol Microbiol 101, 265–280.

17. Monteiro, JM et al. (2018). Peptidoglycan synthesis drives an FtsZ-treadmilling-independent step of cytokinesis. Nature 554, 528–532.

18. Ovesný, M, Křížek, P, Borkovec, J, Švindrych, Z, and Hagen, GM (2014). ThunderSTORM: a comprehensive ImageJ plug-in for PALM and STORM data analysis and super-resolution imaging. Bioinformatics 30, 2389–2390.

19. Perez, AJ et al. (2019). Movement dynamics of divisome proteins and PBP2x:FtsW in cells of *Streptococcus pneumoniae*. Proc Natl Acad Sci U S A 116, 3211–3220.

20. Pichoff, S, and Lutkenhaus, J (2005). Tethering the Z ring to the membrane through a conserved membrane targeting sequence in FtsA. Mol Microbiol 55, 1722–1734.

21. Pichoff, S, and Lutkenhaus, J (2007). Identification of a region of FtsA required for interaction with FtsZ. Mol Microbiol 64, 1129–1138.

22. RayChaudhuri, D, and Park, JT (1992). Escherichia coli cell-division gene ftsZ encodes a novel GTP- binding protein. Nature 359, 251–254.

23. Schindelin, J et al. (2012). Fiji: an open-source platform for biological-image analysis. Nat Methods 9, 676–682.

24. Schoenemann, KM, Krupka, M, Rowlett, VW, Distelhorst, SL, Hu, B, and Margolin, W (2018). Gain-of- function variants of FtsA form diverse oligomeric structures on lipids and enhance FtsZ protofilament bundling. Mol Microbiol 109, 676–693.

25. Schrader, JM, and Shapiro, L (2015). Synchronization of *Caulobacter crescentus* for Investigation of the Bacterial Cell Cycle. Journal of Visualized Experiments.

26. Squyres, GR, Holmes, MJ, Barger, SR, Pennycook, BR, Ryan, J, Yan, VT, and Garner, EC (2021). Single- molecule imaging reveals that Z-ring condensation is essential for cell division in Bacillus subtilis. Nat Microbiol 6, 553–562.

27. Stricker, J, and Erickson, HP (2003). In Vivo Characterization of *Escherichia coli ftsZ* Mutants: Effects on Z-Ring Structure and Function. J Bacteriol 185, 4796–4805.

28. Sundararajan, K, and Goley, ED (2017). The intrinsically disordered C-terminal linker of FtsZ regulates protofilament dynamics and superstructure in vitro. J Biol Chem 292, 20509–20527.

29. Sundararajan, K, Miguel, A, Desmarais, SM, Meier, EL, Casey Huang, K, and Goley, ED (2015). The bacterial tubulin FtsZ requires its intrinsically disordered linker to direct robust cell wall construction. Nat Commun 6, 7281.

30. Sundararajan, K, Vecchiarelli, A, Mizuuchi, K, and Goley, ED (2018). Species- and C-terminal linker- dependent variations in the dynamic behavior of FtsZ on membranes *in vitro*. Mol Microbiol 110, 47–63.

31. Szwedziak, P, Wang, Q, Freund, SM V, and Löwe, J (2012). FtsA forms actin-like protofilaments. EMBO J 31, 2249–2260.

32. Thanbichler, M, and Shapiro, L (2006). MipZ, a Spatial Regulator Coordinating Chromosome Segregation with Cell Division in *Caulobacter*. Cell 126, 147–162.

33. Wang, Y, Jones, BD, and Brun, Y V. (2001). A set of *ftsZ* mutants blocked at different stages of cell division in *Caulobacter*. Mol Microbiol 40, 347–360.

34. Woldemeskel, SA et al. (2020). The conserved transcriptional regulator CdnL is required for metabolic homeostasis and morphogenesis in *Caulobacter*. PLoS Genet 16, e1008591.

35. Woldemeskel, SA, McQuillen, R, Hessel, AM, Xiao, J, and Goley, ED (2017). A conserved coiled-coil protein pair focuses the cytokinetic Z-ring in *Caulobacter crescentus*. Mol Microbiol 105, 721–740.

36. Yang, X et al. (2021). A two-track model for the spatiotemporal coordination of bacterial septal cell wall synthesis revealed by single-molecule imaging of FtsW. Nat Microbiol 6, 584–593.

37. Yang, X, Lyu, Z, Miguel, A, McQuillen, R, Huang, KC, and Xiao, J (2017). GTPase activity-coupled treadmilling of the bacterial tubulin FtsZ organizes septal cell wall synthesis. Science (1979) 355, 744–747.

