## Supplemental Information for "Intrinsic and extrinsic factors regulate FtsZ function in *Caulobacter crescentus*"

Figure S1

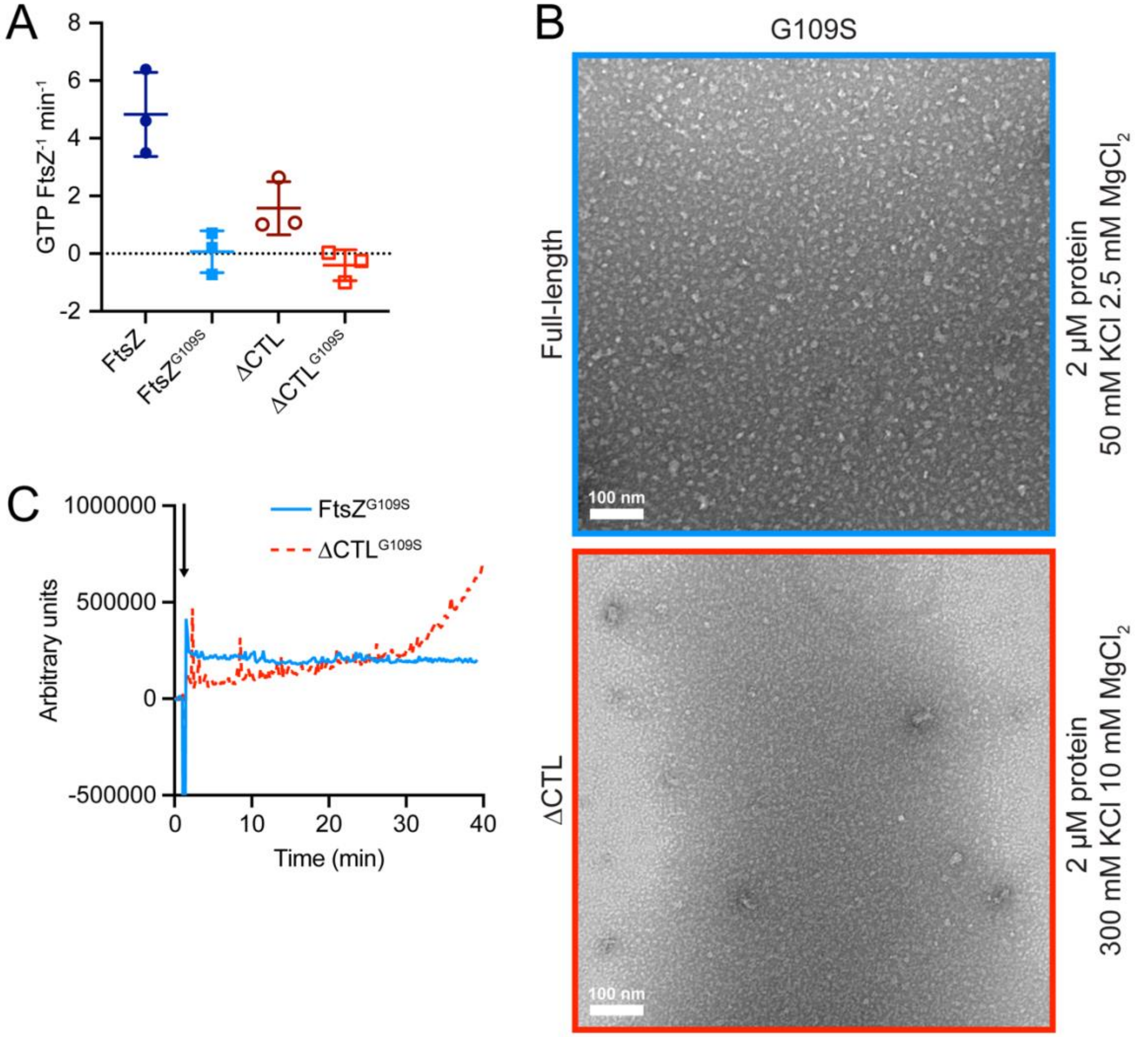


**Supplemental Figure 1: G109S variants fail to form polymers *in vitro***

**A.** GTP turnover rates for indicated FtsZ variants (4 μM). **B.** Representative TEM micrographs for 2 μM of each indicated protein incubated with 2 mM GTP. **C.** Right-angle light scattering at 350 nm over time for 4 μM of each indicated protein upon addition of 0.5 mM GTP, indicated by the black arrow. Representative lines of two independent replicates are shown.

### Figure S2

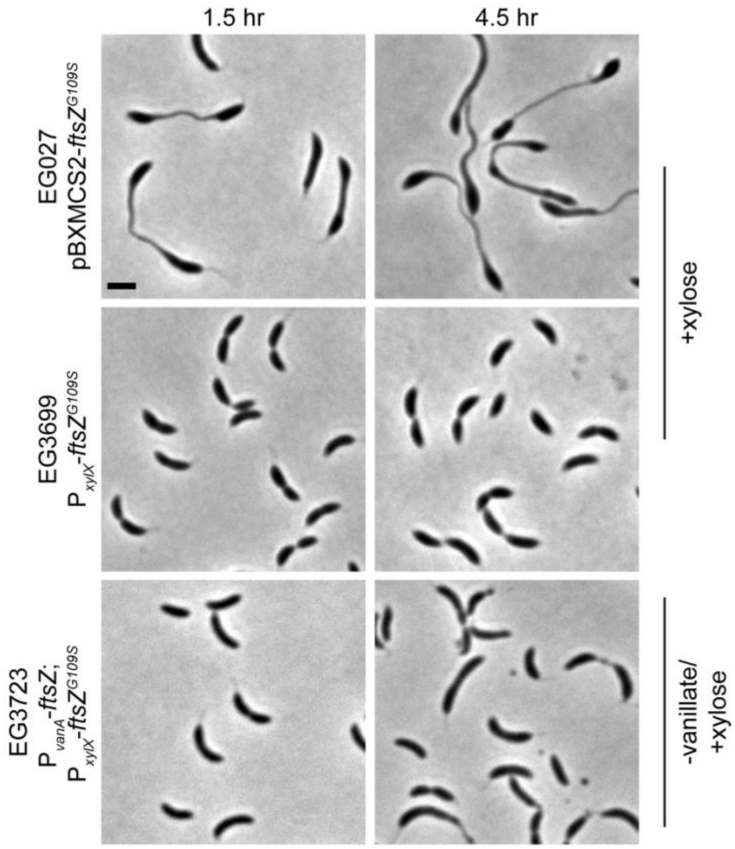


**Supplemental Figure 2: *ftsZ$^{G109S}$* "dumbbell" phenotype is a result of overexpression**

Representative phase contrast micrographs of indicated strains either induced with 0.3% xylose or depleted of vanillate followed by induction with 0.3% xylose for indicated times prior to imaging. Scale bar, 2 µm.

### Figure S3

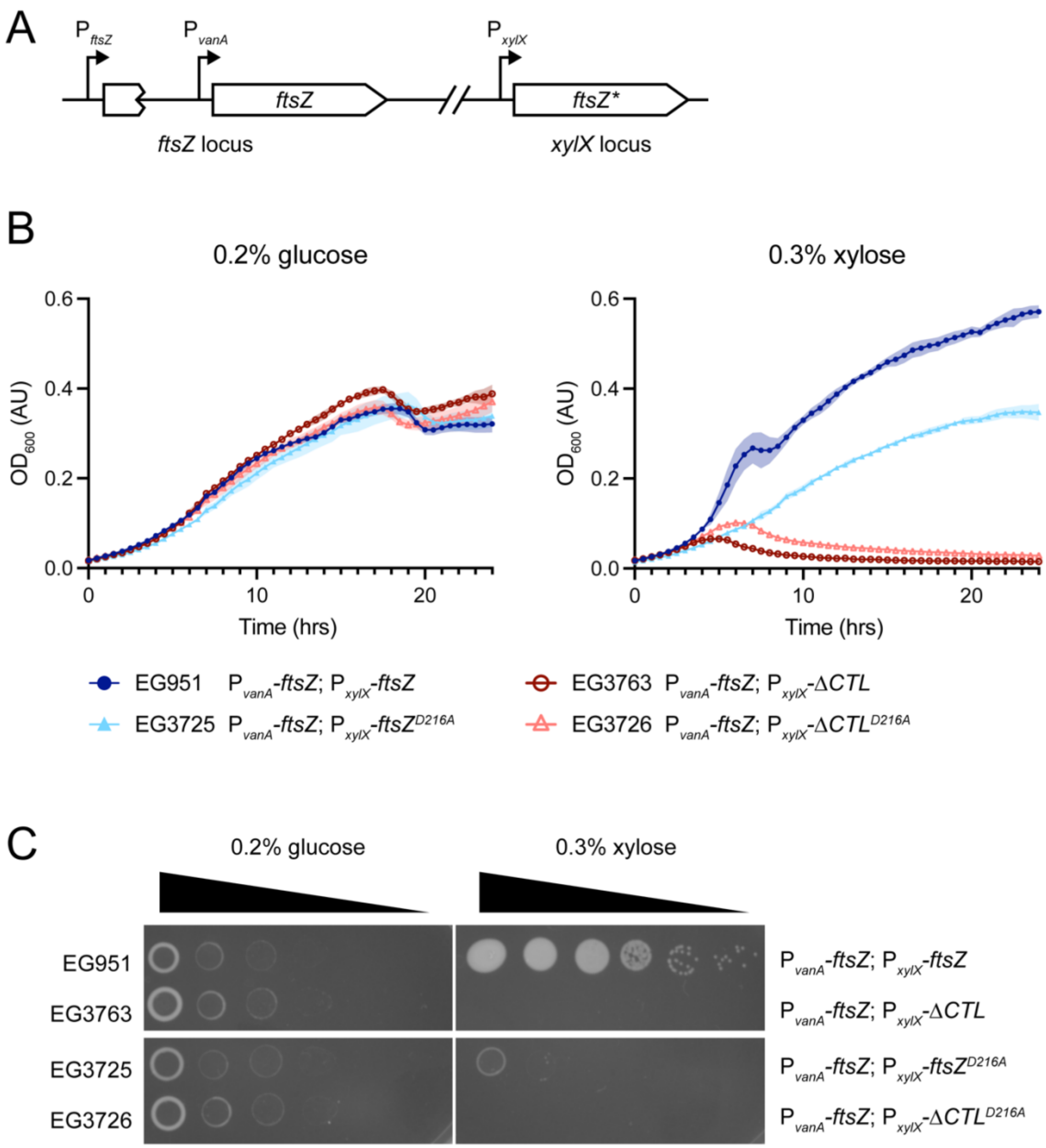


**Supplemental Figure 3: GTPase-deficient FtsZ variants exhibit impaired growth and viability defects**

**A.** Schematic of strategy for simultaneously depleting wild-type FtsZ, under control of the $P_{vanA}$ promoter, while inducing expression of the desired *ftsZ* mutant (*ftsZ**), under control of the $P_{xylX}$ promoter. **B.** Growth curves of indicated strains following simultaneous depletion of vanillate and induction of xylose-driven expression of indicated *ftsZ* with 0.2% glucose (left, control) or 0.3% xylose (right). Lines are mean values of 3 technical replicates (SD shown with shaded regions). **C.** 10-fold serial dilutions of indicated strains plated onto PYE with either 0.2% glucose or 0.3% xylose.

### Figure S4

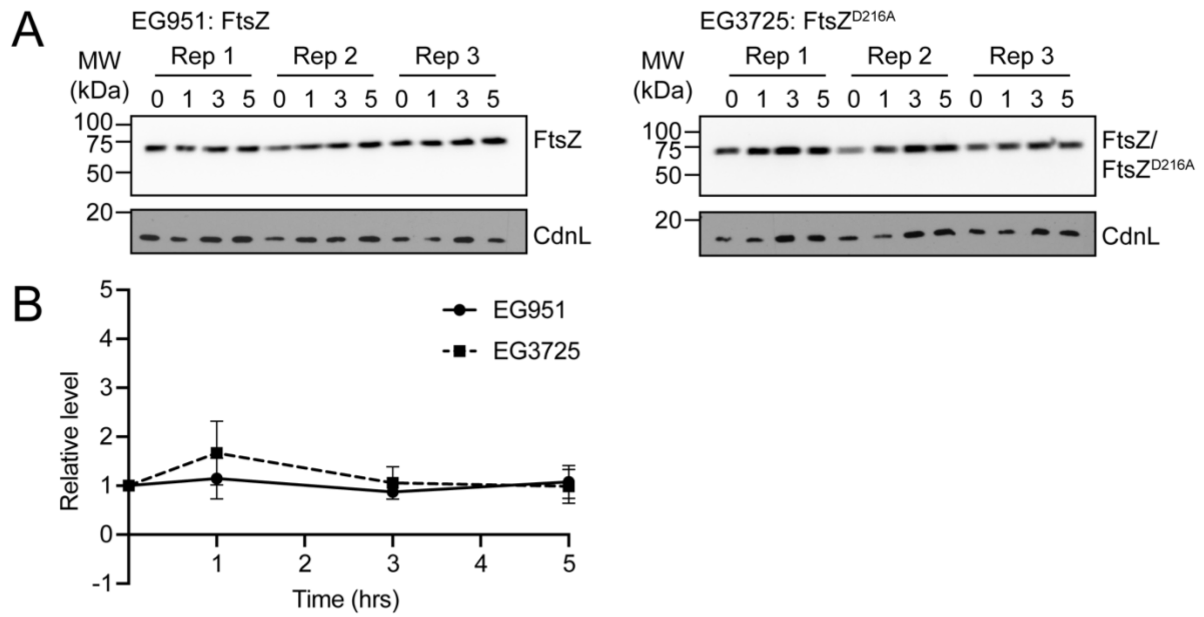


**Supplemental Figure 4: GTPase activity does not affect FtsZ stability *in vivo***

**A.** Immunoblots of indicated strains bearing indicated xylose-inducible FtsZ mutants induced for indicated amounts of time. Each experiment was performed in biological triplicate. Blots are probed with $\alpha$-FtsZ (top) and $\alpha$-CdnL (bottom, loading control) primary antibodies. **B.** Quantification of protein levels in blots from **(A).** FtsZ band intensity was normalized to corresponding loading control band intensity and then normalized to the 0-hr time point for each replicate. Error bars represent standard deviation.

#### Figure S5

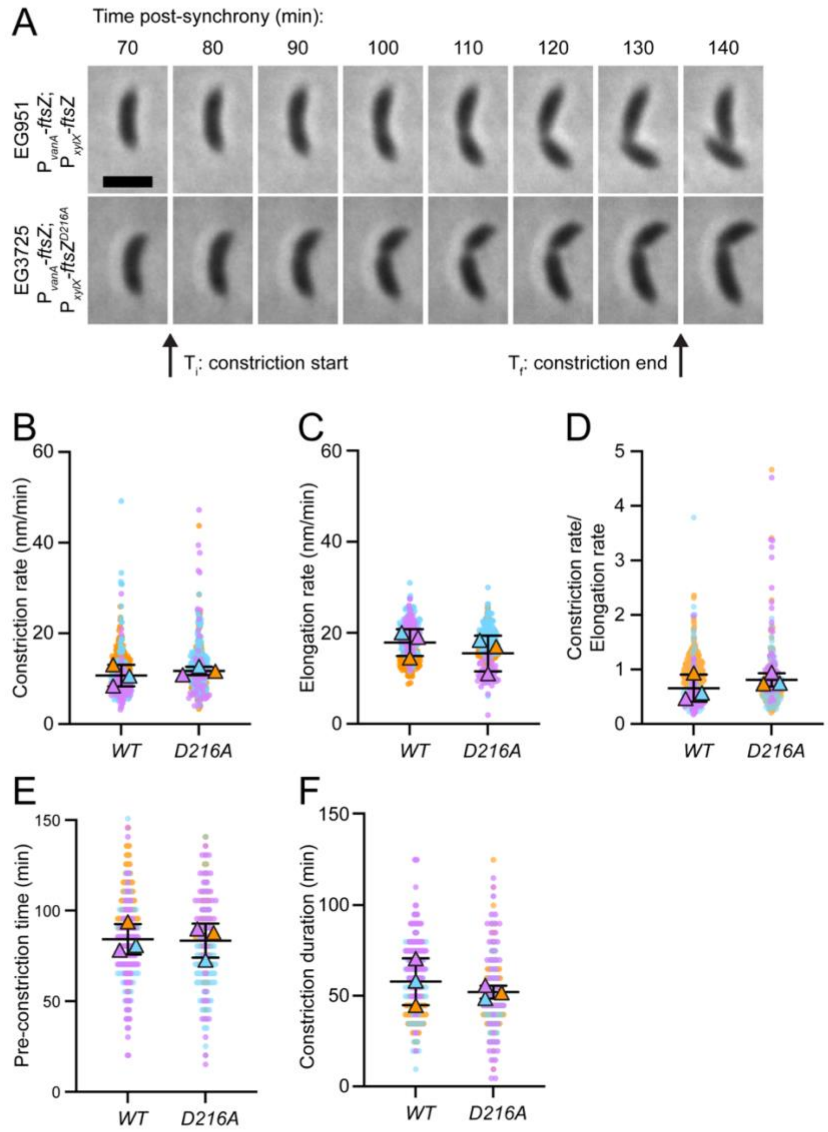


**Supplemental Figure 5: GTPase activity alone does not affect kinetics of constriction**

**A.** Montage of representative phase micrographs of a cells from indicated strains at indicated time points following synchrony. Constriction start ($T_i$) and constriction end ($T_f$) times are indicated below images. Scale bar, 2 µm. **B.-F.** Dot plots of constriction rate (**B**), elongation rate (**C**), the ratio of constriction rate to elongation rate (**D**), pre-constriction time (**E**), or constriction duration (**F**) for strains inducible for the indicate FtsZ mutant. Circles represent individual cell measurements from three independent replicates (orange, cyan, and magenta) and outlined triangles represent mean values for each replicate. Line indicates mean of replicate means and error bars are standard deviation. A parametric student's t-test was performed to compare values for each measurement, none of which were determined to be significantly different.

### Figure S6

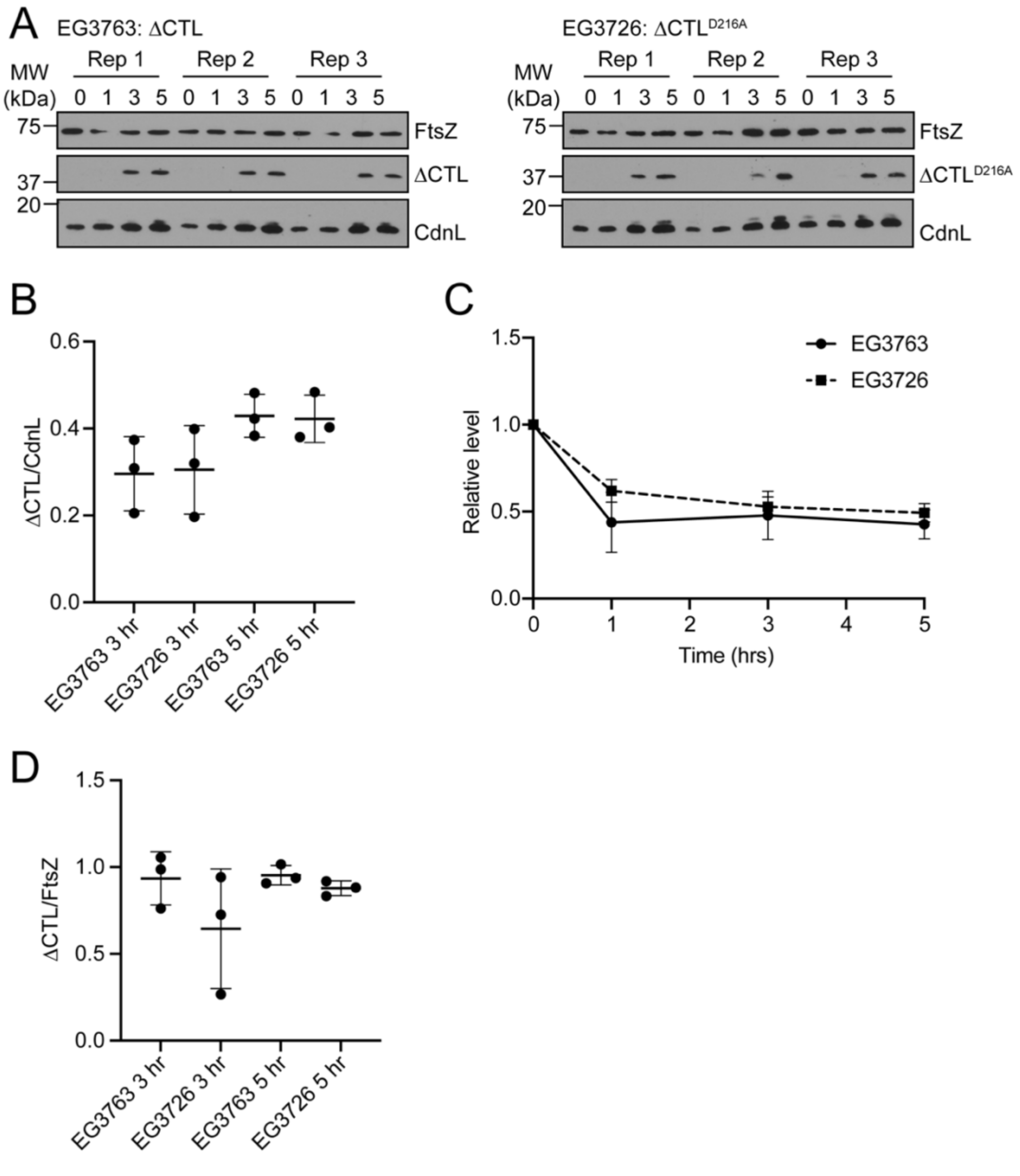


**Supplemental Figure 6: GTPase activity does not affect ΔCTL stability *in vivo***

**A.** Immunoblots of indicated strains bearing indicated xylose-inducible ΔCTL mutants induced for indicated amounts of time. Each experiment was performed in biological triplicate. Blots are probed with α-FtsZ (top two) and α-CdnL (bottom, loading control) primary antibodies. **B.** Quantification of ratios of ΔCTLvariant/CdnL protein levels in blots from (**A**) at 3- and 5-hr time points. **C.** Quantification of wild-type FtsZ protein levels in blots from (**A**). Wild-type FtsZ band intensity was normalized to corresponding loading control band intensity and then normalized to the 0-hr time point for each replicate. **D.** Ratios of indicated ΔCTL variant/FtsZ protein levels in blots from (**A**) at 3- and 5-hr time points. Error bars represent standard deviation.

### Figure S7

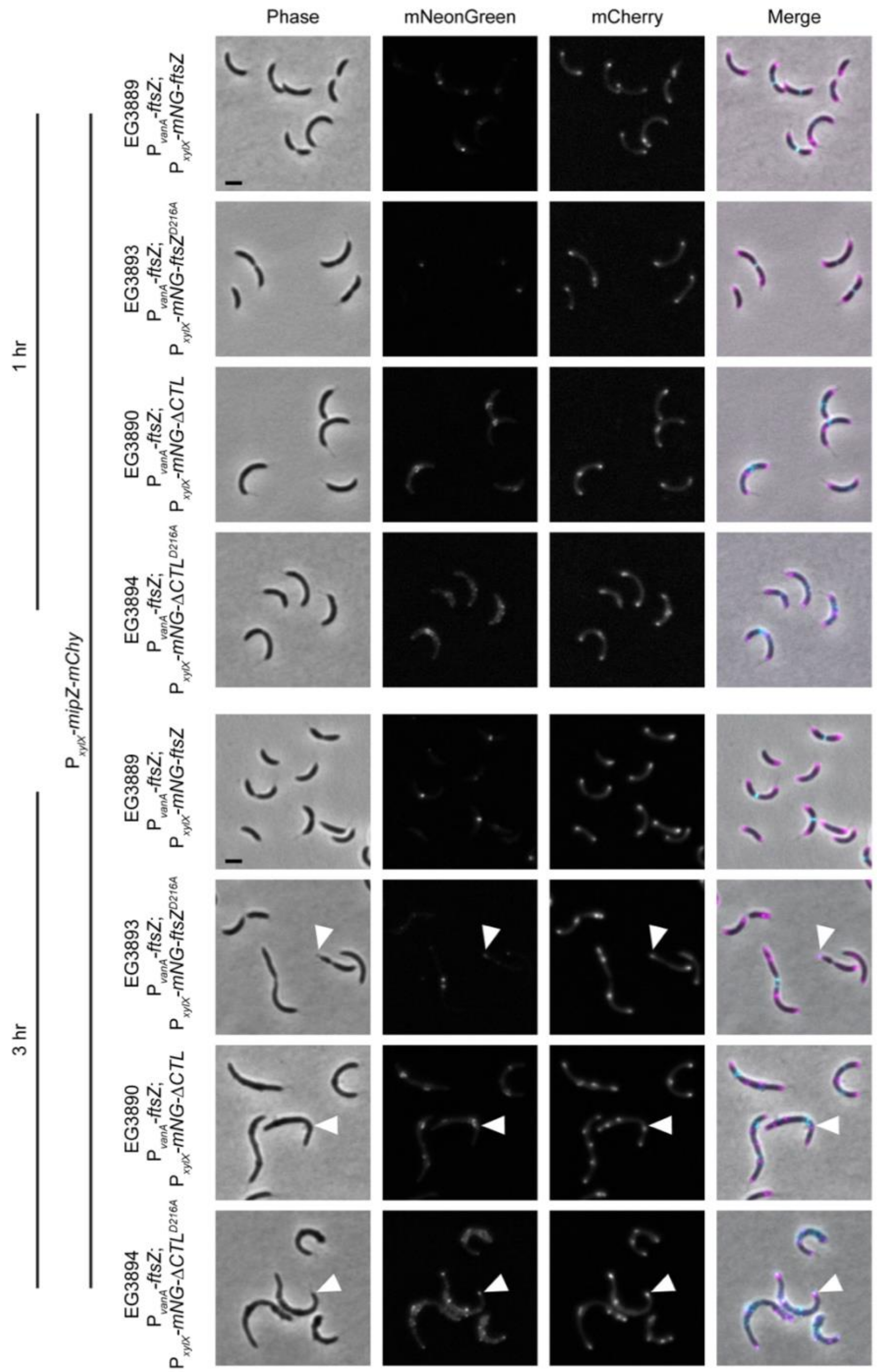


**Supplemental Figure 7: MipZ exhibits normal localization in the presence of GTPase deficient FtsZ variants**

Representative phase contrast, epifluorescence, and merged micrographs of indicated strains at given time points following simultaneous depletion of vanillate and induction of xylose-driven expression of indicated *mNeonGreen-ftsZ* fusion and *mipZ-mCherry* with 0.3% xylose. mNeonGreen and mCherry are represented in cyan and magenta, respectively, in the merged image. White arrowheads indicate instances where the indicated mNG-FtsZ variant and MipZ-mCherry are close together. Scale bar, 2 μm.

### Figure S8

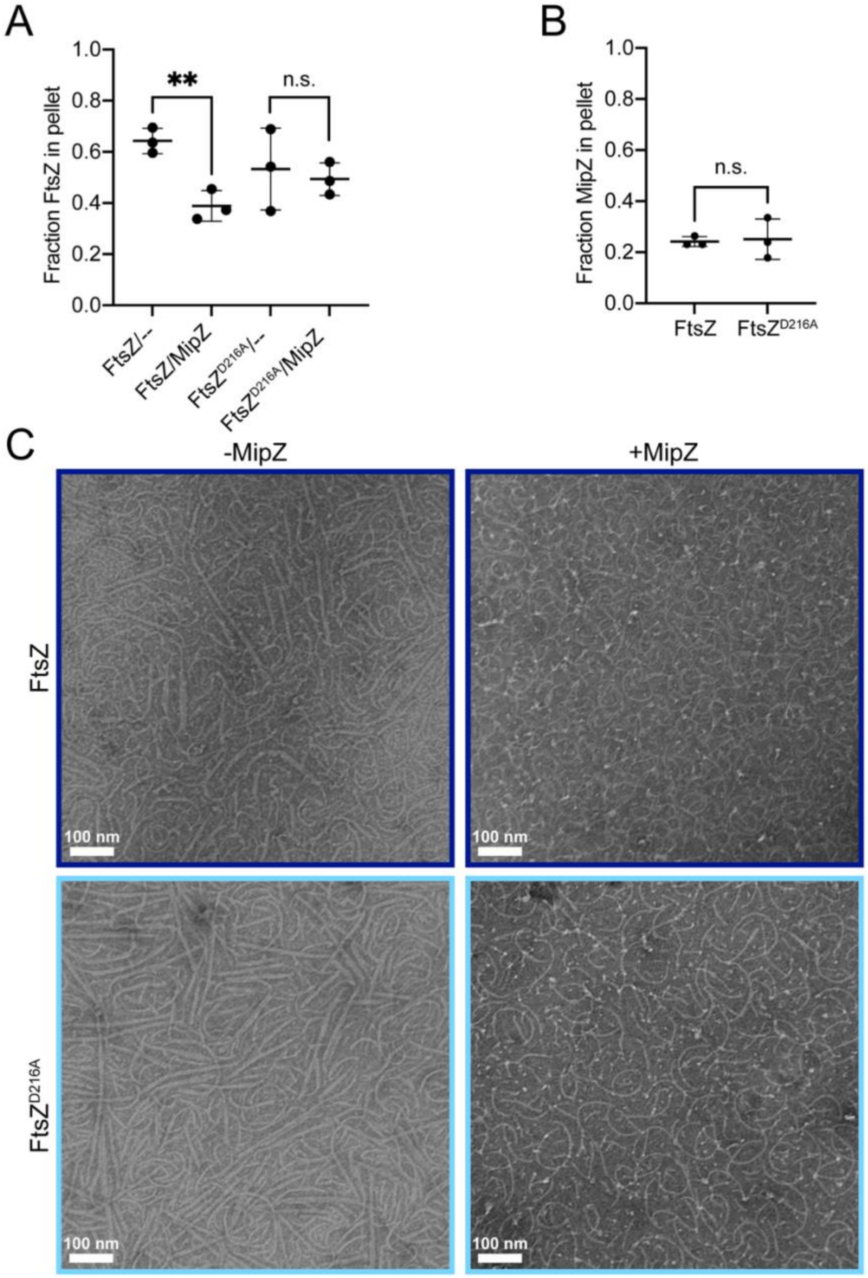


**Supplemental Figure 8: FtsZ GTPase activity modulates regulation by MipZ**

**A.- B.** Quantification of fraction of FtsZ (3 µM) (**A**) or MipZ (6 µM) (**B**) in pellet upon centrifugation following polymerization with the indicated protein components with 2 mM GTP/2 mM ATP. Experiments were completed in triplicate and parametric student's t-tests were performed on indicated pairs of columns to determine indicated *p* values (n.s., not significant; **, $p \leq 0.01$). **C.** Representative TEM micrographs for 3 µM indicated FtsZ variant incubated with 2 mM GTP/2 mM ATP without or with MipZ (6 µM). Scale bar, 100 nm.

### Figure S9

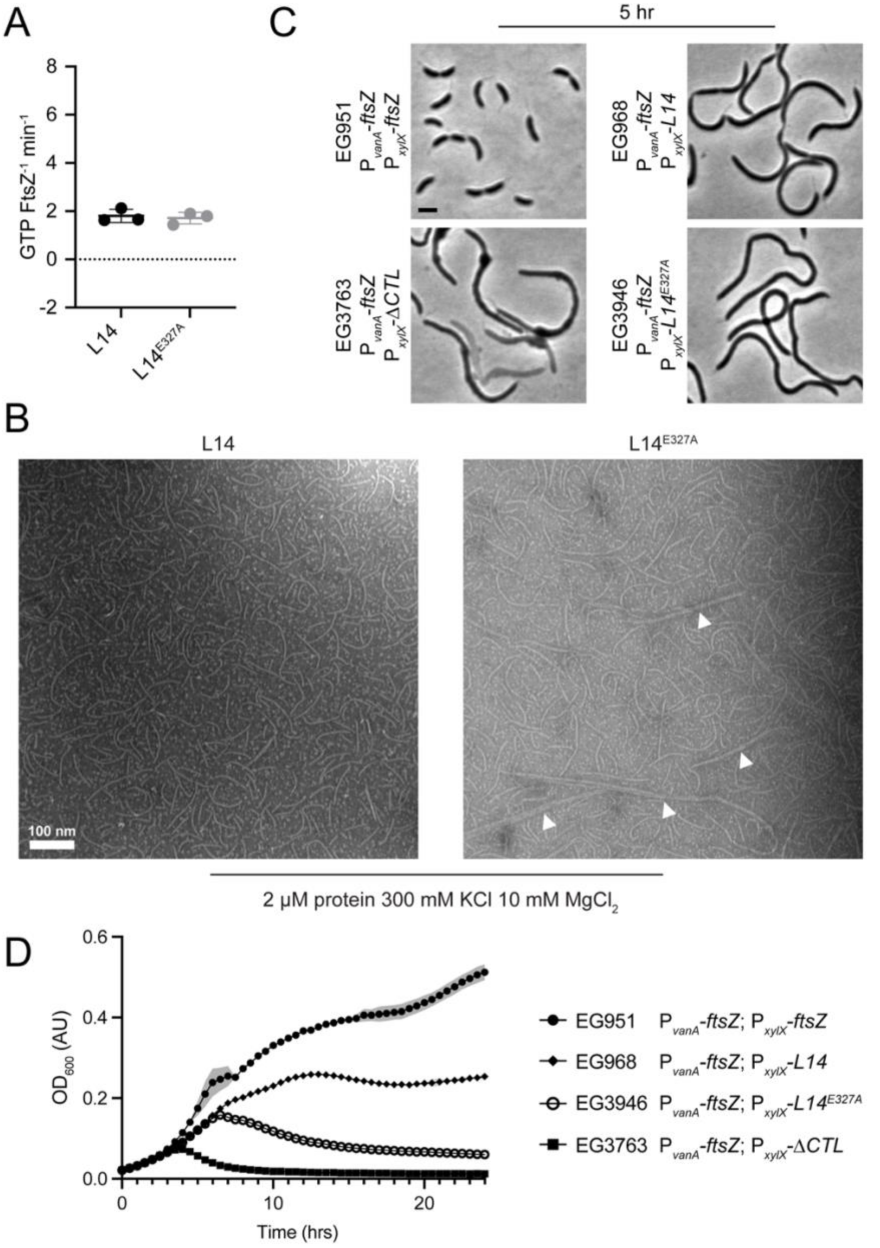


**Supplemental Figure 9: Loss of both charge and length are required for bundling, bulge formation, and lysis characteristic of ΔCTL**

**A.** GTP turnover rates for indicated FtsZ variants (4 mM). **B.** Representative TEM micrographs for 2 μM of each indicated protein incubated with 2 mM GTP. Bundles formed by L14 E327A are indicated by white arrowheads. **C.** Representative phase contrast micrographs of indicated strains at given time points following simultaneous depletion of vanillate and induction of xylose-driven expression of indicated *ftsZ* with 0.3% xylose. Scale bar, 2 μm. **D.** Growth curves of indicated strains following simultaneous depletion of vanillate and induction of xylose-driven expression of indicated *ftsZ* with 0.3% xylose. Lines are mean values of 3 technical replicates (SD shown with shaded regions).

**Supplemental Table 1:** A list of *Caulobacter crescentus* strains used in this study.

**Supplemental Table 2:** A list of plasmids used in this study.
